# Concentration-dependent formation of intersegment interactions in viral droplets of influenza A virus infected cells

**DOI:** 10.1101/2024.09.29.615319

**Authors:** Naoki Takizawa, Koichi Higashi, Risa Karakida Kawaguchi, Yasuhiro Gotoh, Tetsuya Hayashi, Ken Kurokawa

**Affiliations:** Laboratory of Virology, Institute of Microbial Chemistry (BIKAKEN), 3-14-23 Kamiosaki, Shinagawa-ku, Tokyo 141-0021, Japan; Genome Evolution Laboratory, National Institute of Genetics, 1111 Yata, Mishima, Shizuoka 411-8540, Japan; Center for iPS Cell Research and Application, Kyoto University, 53 Kawahara-cho, Shogoin, Sakyo-ku, Kyoto 606-8507, Japan; Department of Bacteriology, Faculty of Medical Sciences, Kyushu University, 3-1-1 Maidashi, Higashi-ku, Fukuoka 812-8582, Japan

## Abstract

The influenza A virus genome consists of eight RNA segments, each incorporated into a virion. It has been proposed that intersegment interactions bundle these segments, and viral inclusions, which contain viral ribonucleoproteins (vRNPs) and Rab11, facilitate this process. However, the locations and mechanisms of intersegment interaction formation remain unclear. To investigate this, we identified comprehensive intersegment interactions in infected cells using customized LIGR-seq. Our results revealed that intersegment interactions in infected cells overlapped with those in the virion and were partially formed through direct RNA-RNA interactions. Additionally, we found that the formation of these interactions was delayed in cells expressing a Rab11 dominant-negative. Furthermore, artificially increasing vRNP concentrations in the nucleus, where intersegment interactions typically do not occur, triggered the formation of these interactions, which again overlapped with those in the virion. These results suggest that intersegment interactions are formed in viral inclusions where progeny vRNPs accumulate.

## Introduction

Influenza A viruses (IAVs) are human and animal respiratory pathogens, and IAV infection is a threat to global health due to significant morbidity and mortality. In addition to seasonal influenza epidemics every year, pandemics caused by IAVs occur every 20-30 years. The IAV genome consists of eight single-stranded negative-sense RNA segments (vRNAs), and the vRNA is complexed with a heterotrimeric viral RNA polymerase complex and nucleoproteins (NP) to form a viral ribonucleoprotein complex (vRNP). vRNP forms a double-helical structure with the polymerase at one end and a short loop at the other ^1^. Pandemic IAVs emerge through genetic reassortment in which human, avian, and/or swine IAVs exchange parts of their segmented genomes ^2^. Genetic reassortment is thought to be intimately linked to the co-packaging of one set of the eight different IAV segmented genomes. However, the molecular mechanisms of how one set of the eight segments is selected and packaged in virion are still poorly understood.

Genomic regions required for IAV replication and genome packaging have been identified. The 12 nucleotides from the 3′ termini and the 13 nucleotides from the 5′ termini of the vRNAs, which are partially complementary and highly conserved between segments, are essential for viral RNA synthesis and packaging ^3^. Segment-specific noncoding regions and viral protein coding regions at the 3′ and 5′ end of each vRNA segment are necessary for efficient viral genome packaging ^4^. Moreover, the extra signal sequences in the middle of coding regions have been found to be required for efficient genome packaging ^5^. Mutations and deletions of the signal sequences resulted in the reduction of packaging not only of the mutated segment itself but also that of the other segments ^6–12^. While the function of these signal sequences remains unknown, these signal sequences are hypothesized to be involved in intersegment vRNA-vRNA interactions. Regions responsible for the intersegment interactions have been identified in vitro, and some intersegment interactions are required for genome packaging ^5,13–15^. Moreover, direct contacts between the vRNPs have also been observed by electron tomographic analyses ^16^. Recently, comprehensive intersegment interaction analysis using high-throughput sequencing (HTS) has been conducted to elucidate the relationship between intersegment interactions and genome packaging ^17–19^. These studies demonstrated a redundant and complex network of intersegment interactions in the virion. Dadonaite et al. showed that intersegment interactions in virion are formed by direct RNA-RNA interactions and drive bundling of eight segments during reassortment between different IAV strains ^17^. However, other studies showed that disruption of intersegment interactions does not affect viral propagation ^18,19^. As such, intersegment interactions are thought to be one of the important factors to control precise genome bundling, but the detailed role is still unknown.

Besides the function of the packaging signal, where the segmented genome assembles in infected cells is important for understanding the mechanism of segment assembly. The replication of vRNA occurs in the nucleus, and newly synthesized vRNPs are exported from the nucleus via a CRM1-dependent pathway ^20,21^. The exported vRNPs are transported to the apical plasma membrane by interacting with Rab11, which is associated with recycling endosomes ^22–24^, and eight segments are assembled in the cytoplasm ^25–27^. Rab11 is a small GTPase belonging to the Ras-related superfamily. Rabs are associated with cyclical activation and inactivation in the form of GTP-bound/active state and GDP-bound/inactive state ^28^. The subcellular localization of Rab11 in infected cells is changed ^29–31^, and Rab11 was distributed in the cytoplasm as an enlarged puncta where multiple segments are colocalized during the late phase of infection ^27,29^. These enlarged vRNP/Rab11 puncta have properties of liquid droplets, forming a condensate in the cell ^32^. There are many viral condensates formed by RNA virus infections. These viral condensates were shown recently to have liquid properties that adopt many names like virus factories, viroplasm, or viral inclusions and are thought to provide a physical platform to concentrate viral genome and proteins for operating in different steps of the viral replication cycle, including replication and assembly ^33,34^. Viral inclusions in IAV infected cells are hypothesized to be a platform to bundle eight segments by intersegment interactions ^32^. However, it is not clear whether intersegment interactions in infected cells are identical to those in the virion.

Here, we identify comprehensive intersegment interactions of IAV in infected cells by applying the HTS approach and show that intersegment interactions overlapping with those in the virion are identified under the conditions of forming viral inclusions. Even with artificially increased vRNP concentrations in the nucleus, intersegment interactions also overlapped with those in the virion. Our results indicate that the intersegment interaction network is formed in a vRNP concentration-dependent manner.

## Results

### Intersegment interactions in purified virion

To identify comprehensive RNA interactions in virion and infected cells, we combined ligation of interacting RNA and high-throughput sequencing (LIGR-seq) with an RNA pull-down procedure and customized data analysis (Figure S1). Specifically, a vRNA purification step using single-stranded cDNA probes for vRNAs is added to the normal LIGR-seq process. To detect intersegment interactions more quantitatively, we have further improved analytical methods for LIGR-seq as follows: 1) The contact map was normalized using the iterative method that was employed in Hi-C data analysis, 2) the normalized count in each 100 nucleotide (nt) bin was referred to as the contact score, then 3) we identified reliable intersegment interactions by adjusting the contact scores from the duplicate experiments using the irreproducible discovery rate (IDR) (see Materials and methods). The number of inter- and intrasegment interaction reads is sufficient for downstream analysis. However, there appears to be a bias in the read distribution (Figure S2). Therefore, we normalized the contact map using both inter- and intrasegment reads. Moreover, since many non-specific intersegment interactions are detected in the LIGR-seq, we used IDR to extract reliable intersegment interactions (Figure S2).

First, we identified comprehensive intersegment interactions in the virion by the customized LIGR-seq. We identified 659 intersegment interactions in the virion (Figures 1A and S2A). A list of intersegment interaction is provided in the supplemental data. As reported previously, intersegment interactions were redundant and complex, and “hotspots” of intersegment interactions were detected ^17,18^. Hotspots of intersegment interactions were predominantly located at the 3′ and 5′ ends and in the middle of the segment (Figure S3). This result suggests that a significant portion of intersegment interactions are formed among 3′ and 5′ ends and the middle of the segment, which may reflect that all eight segments are not packaged in a virion in a head (polymerase side) - to-tail (loop side) orientation ^35^. We performed customized LIGR-seq using purified vRNA samples to assess non-specific intersegment interactions in customized LIGR-seq, and 235 intersegment interactions were identified (Figures 1B and S2B). Among the 235 intersegment interactions identified in purified vRNA, 21 were reproduced at the same locations in the virion. Since there is a mismatch between the intersegment interaction site and the RNA ligation site in LIGR-seq, we also quantified the number of intersegment interactions overlapping within 100 nt in the virion and vRNA. Among the 235 intersegment interactions, 78 were reproduced within 100 nt in the virion. To test the significance of intersegment interaction overlaps in the virion and purified vRNA, the expected number of overlapping intersegment interactions is calculated by randomly generating 659 and 235 intersegment interactions. In our random simulation, average numbers of 18.3 and 84.0 intersegment interactions were reproduced at the same locations and within 100 nt in two randomly generated intersegment interaction sets, respectively. Since the calculated p-value is more than 0.05 (*p* = 0.210 and 0.772, respectively) (Figure 1B and Table 1), we conclude that the intersegment interactions identified in vRNA do not significantly overlap with those in the virion.

**Figure 1.**
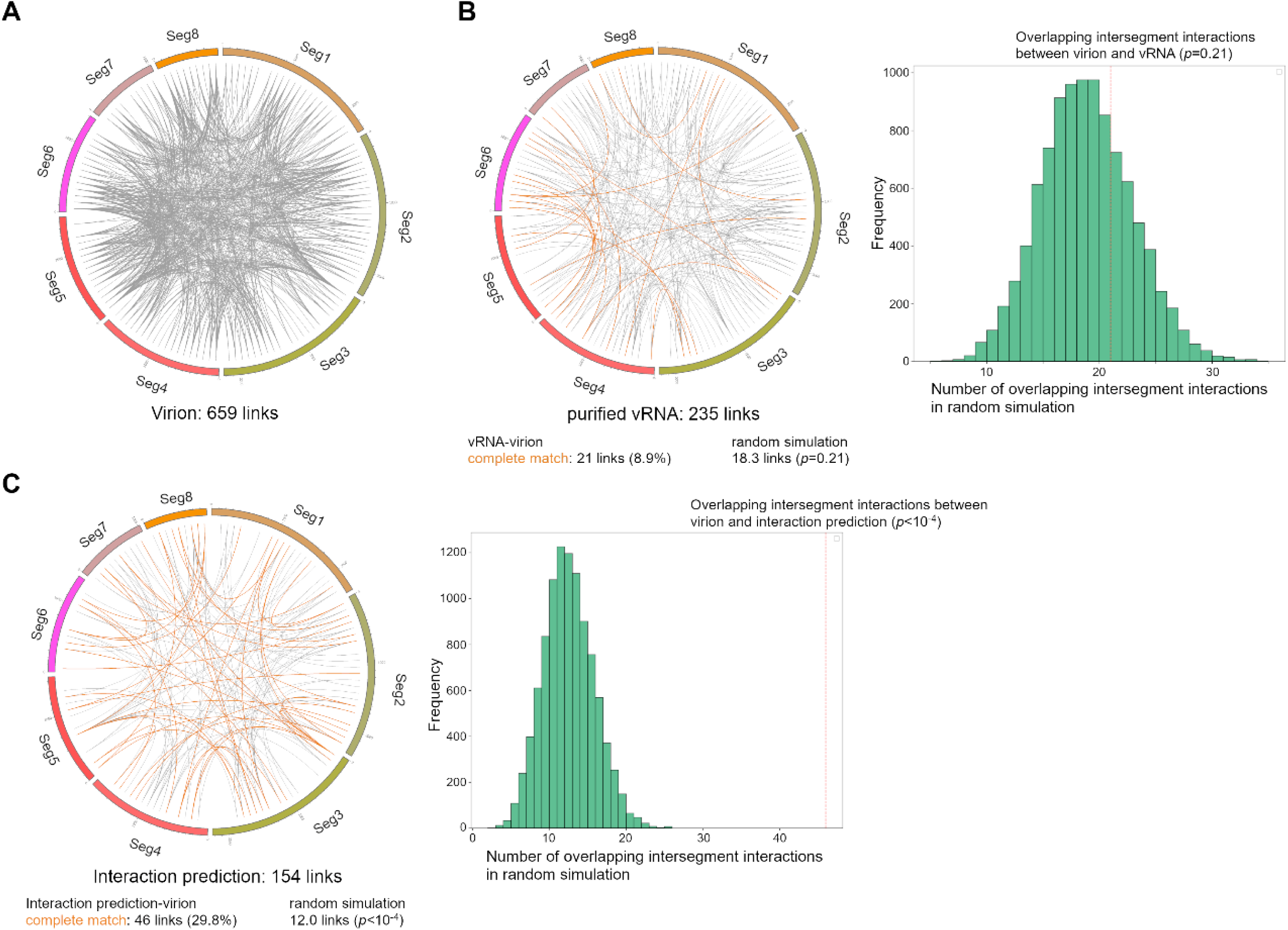
Intersegment interactions in purified virion identified by customized LIGR-seq. (A) Circos plots of the intersegment interactions in the wild type PR8 virus. Each line indicates the intersegment interaction which results from LIGR-seq. (B) Circos plots of the intersegment interactions in purified vRNA. Intersegment interactions identified in both the virion and purified vRNA are indicated by orange lines. “Random simulation” means the number of overlapping intersegment interactions between two randomly generated intersegment interaction sets. The right panel shows the distribution of the number of overlapping intersegment interactions in random simulation. The number of overlapping intersegment interactions between virion and vRNA is indicated by the red dashed line. (C) Circos plots of the predicted intersegment interactions. Intersegment interactions identified in both the virion and the prediction are indicated by orange lines. The right panel shows the distribution of the number of overlapping intersegment interactions in random simulation. The number of overlapping intersegment interactions between virion and interaction prediction is indicated by the red dashed line.

**Table 1.**
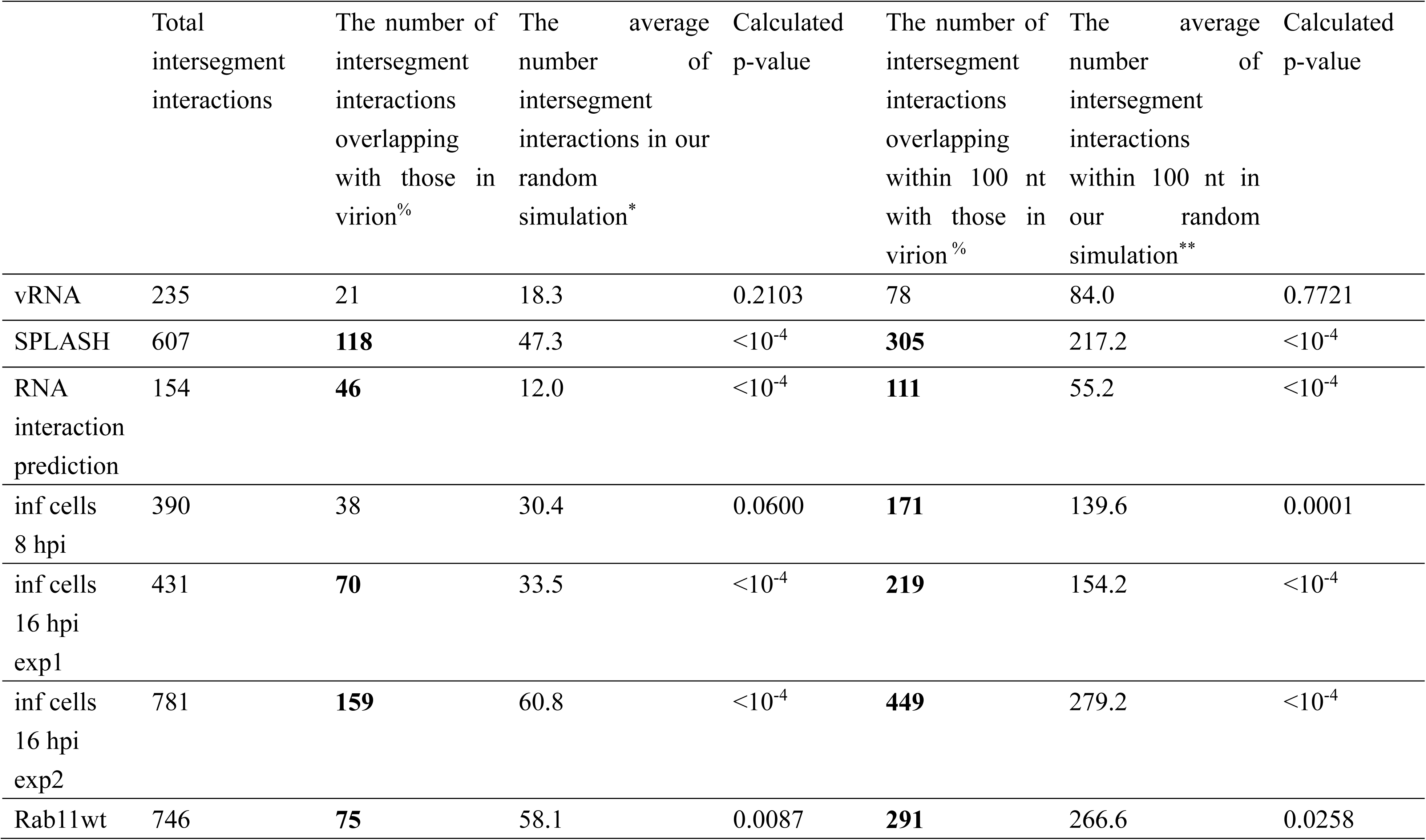
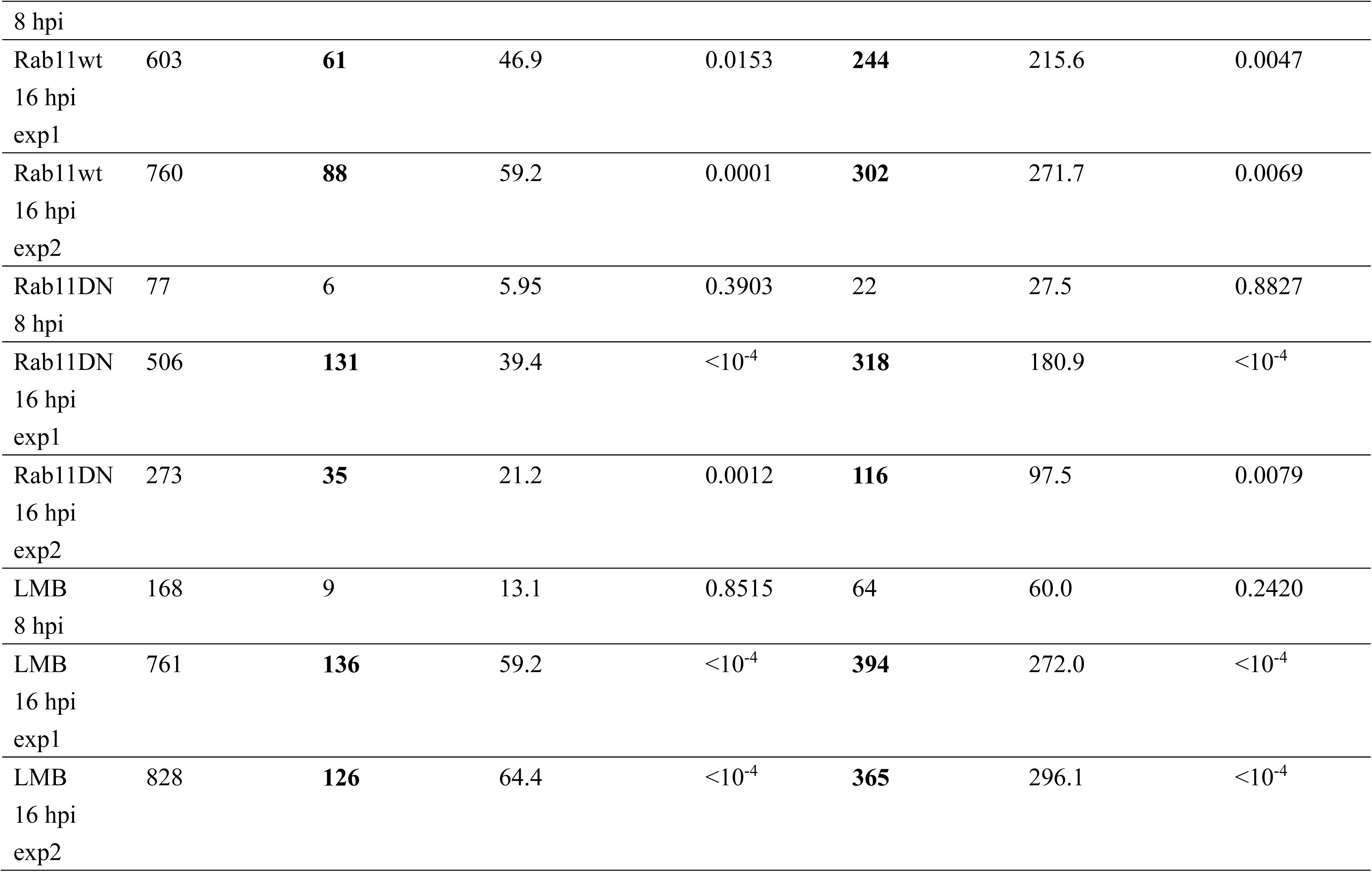
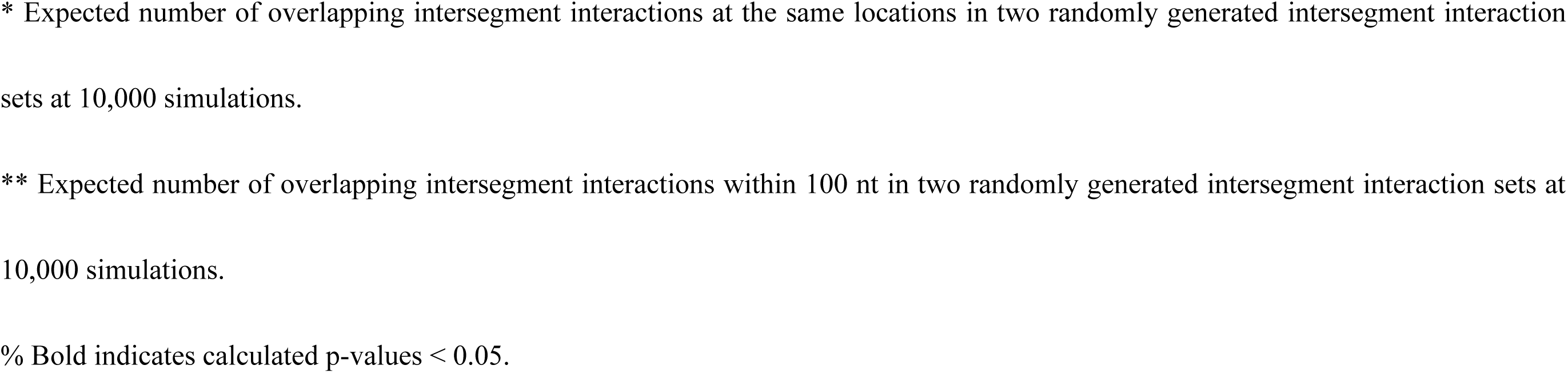
Comparison with intersegment interactions in virion.

Moreover, to confirm the reproducibility of the intersegment interactions captured by our LIGR-seq, we compared identified intersegment interactions in the virion with those previously identified by sequencing of psoralen-crosslinked, ligated, and selected hybrids (SPLASH) ^17^. Among the 607 intersegment interactions identified by SPLASH, 118 and 305 were reproduced at the same locations and within 100 nt in our LIGR-seq result, respectively (Figure S4). In our random simulation, average numbers of 47.3 and 217.2 intersegment interactions were reproduced at the same locations and within 100 nt, respectively, and the calculated p-value was less than 0.05 (*p* < 10^-4^) (Table 1). These results suggest that intersegment interactions in the virion identified by our LIGR-seq significantly overlap with those identified by the previous study.

Next, we examined whether the intersegment interactions identified by customized LIGR-seq were compatible with the prediction of RNA-RNA interactions. The intersegment interactions were predicted by RIblast, which can efficiently predict the pair of regions that are involved in RNA-RNA interactions based on the sequences in the database of RNAs ^36^. Based on the RIblast prediction, 154 intersegment interactions were predicted (Figure 1C). Among the 154 predicted intersegment interactions, 46 and 111 were reproduced at the same locations and within 100 nt in the virion, respectively. In our random simulation, average numbers of 12.0 and 55.2 intersegment interactions were reproduced at the same locations and within 100 nt, respectively, and the calculated p-value was less than 0.05 (*p* < 10^-4^) (Figure 1C and Table 2). These results suggest that the intersegment interactions in the virion are formed mainly by direct RNA interactions through complementary bases. In contrast, the intersegment interactions identified in vRNA do not significantly overlap with the predicted interactions (Table 2). This lack of overlap is likely related to the concentration of the vRNAs.

**Table 2.**
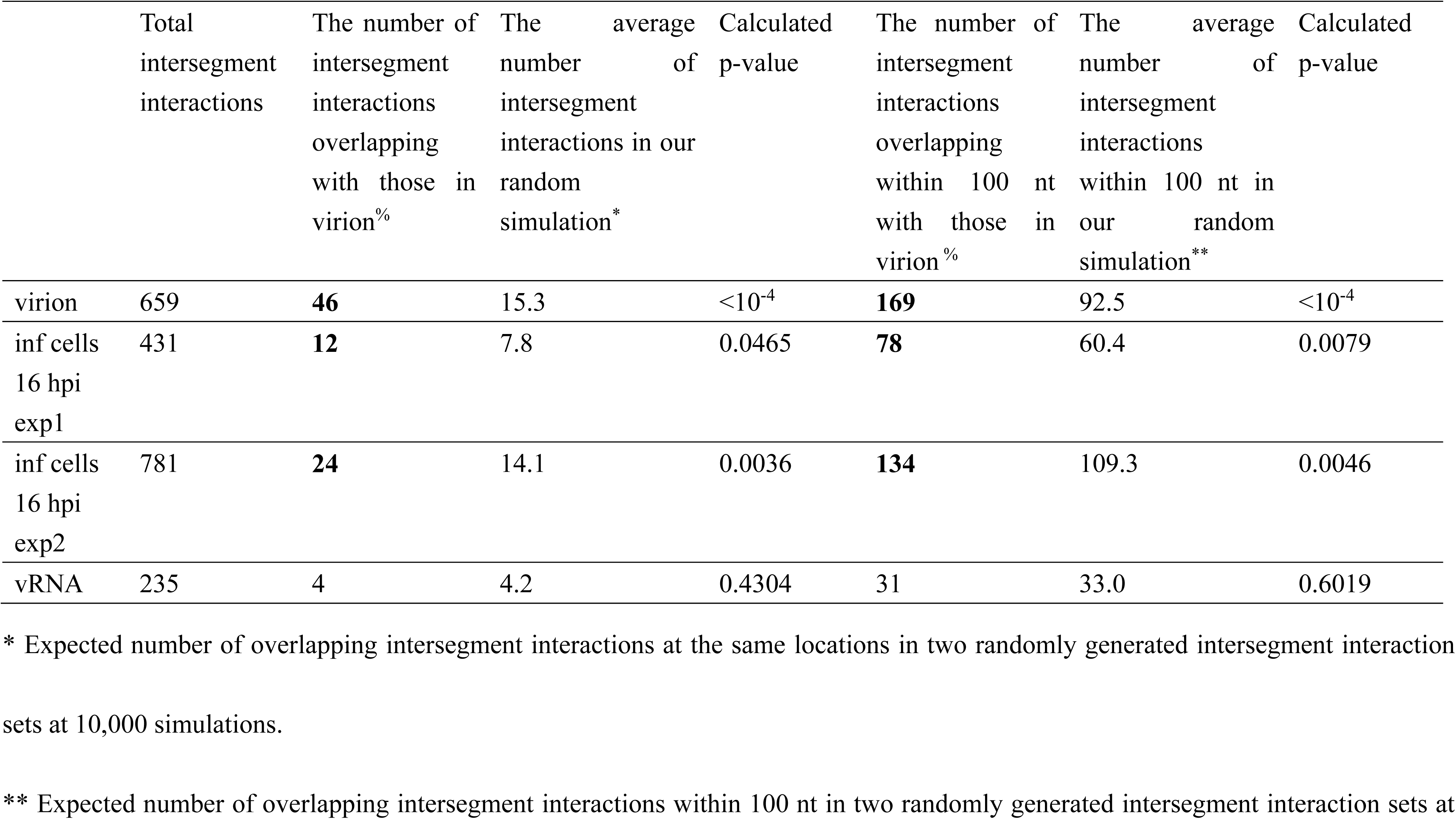
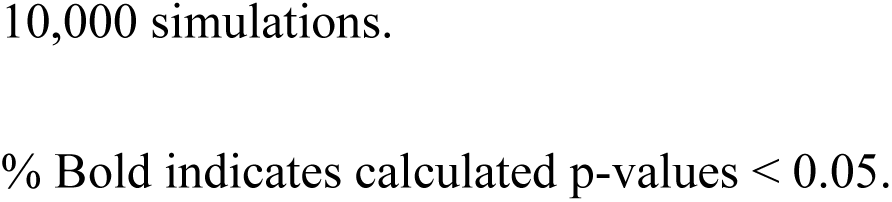
Comparison with predicted intersegment interactions.

### Intersegment interaction in infected cells

Since we successfully analyzed intersegment interactions for the virion via LIGR-seq, we next applied this strategy to uncover the difference of comprehensive intersegment interactions in infected cell conditions. MDCK cells were infected with IAV, and AMT cross-link was performed at 8 and 16 hours post-infection (hpi). In contrast to the virion samples, strong intersegment interaction signals were detected in the infected cell samples both with and without RNA ligation (Figure S5). These signals are due to intersegment recombination of the viral genome during the replication process. To exclude signals caused by recombination, the value of bin, where the signal was detected in the sample without RNA ligation, was set to zero. We identified 390 and 431 intersegment interactions on the IAV genome in infected cells at 8 and 16 hpi, respectively (Figures 2A, 2B, S6). Among them, 38 and 70 intersegment interactions at 8 and 16 hpi were reproduced at the same locations in the virion. In our random simulation, average numbers of 30.4 and 33.5 intersegment interactions were reproduced at the same locations, respectively. The overlapping intersegment interactions between infected cells at 8 hpi and virion are not significantly enriched (*p* = 0.06), whereas those at 16 hpi are significantly enriched (*p* < 10^-4^) (Figures 2B, 2C, and Table 1). Moreover, hotspots of intersegment interactions were frequently observed at the 3′ and 5′ ends and in the middle of the segment in infected cells at 16 hpi (Figure S3). When considering the proximal intersegment interactions within 100 nt, 171 and 219 intersegment interactions at 8 and 16 hpi overlapped with those in the virion. In our random simulation, the average numbers of intersegment interactions reproduced within 100 nt were 139.6 and 154.2, respectively. In this case, we detected significant enrichment in both 8 and 16 hpi conditions (*p* < 10^-4^) (Table 1). The overlapping intersegment interactions between infected cells at 8 hpi and 16 hpi are significantly enriched (*p* = 0.04) (Table 3). We performed two independent experiments on infected cells at 16 hpi, and similar results were obtained (Table 1). These results suggest that intersegment interactions in infected cells at 16 hpi overlapped with those in the virion, even though weakly at 8 hpi.

**Figure 2.**
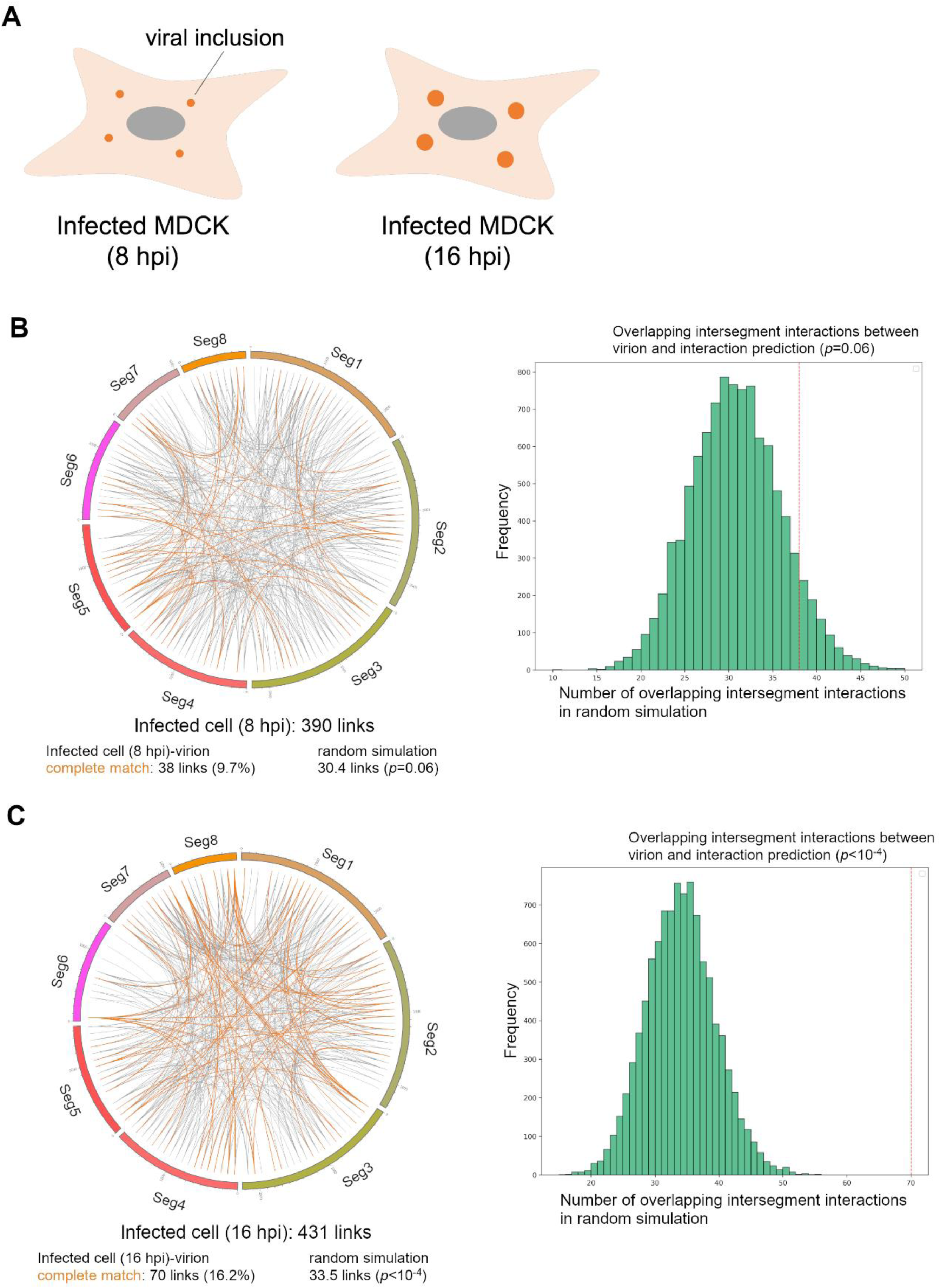
Intersegment interactions in infected cells. (A) Representative depiction of cells subjected to LIGR-seq. (B and C) Circos plots of the intersegment interactions in MDCK cells at 8 hpi (B) and 16 hpi (C). Intersegment interactions identified in both the virion and the cell are indicated by orange lines. “Random simulation” means the number of overlapping intersegment interactions between two randomly generated intersegment interaction sets. The right panel shows the distribution of the number of overlapping intersegment interactions in random simulation. The number of overlapping intersegment interactions between virion and infected cells is indicated by the red dashed line.

**Table 3.**
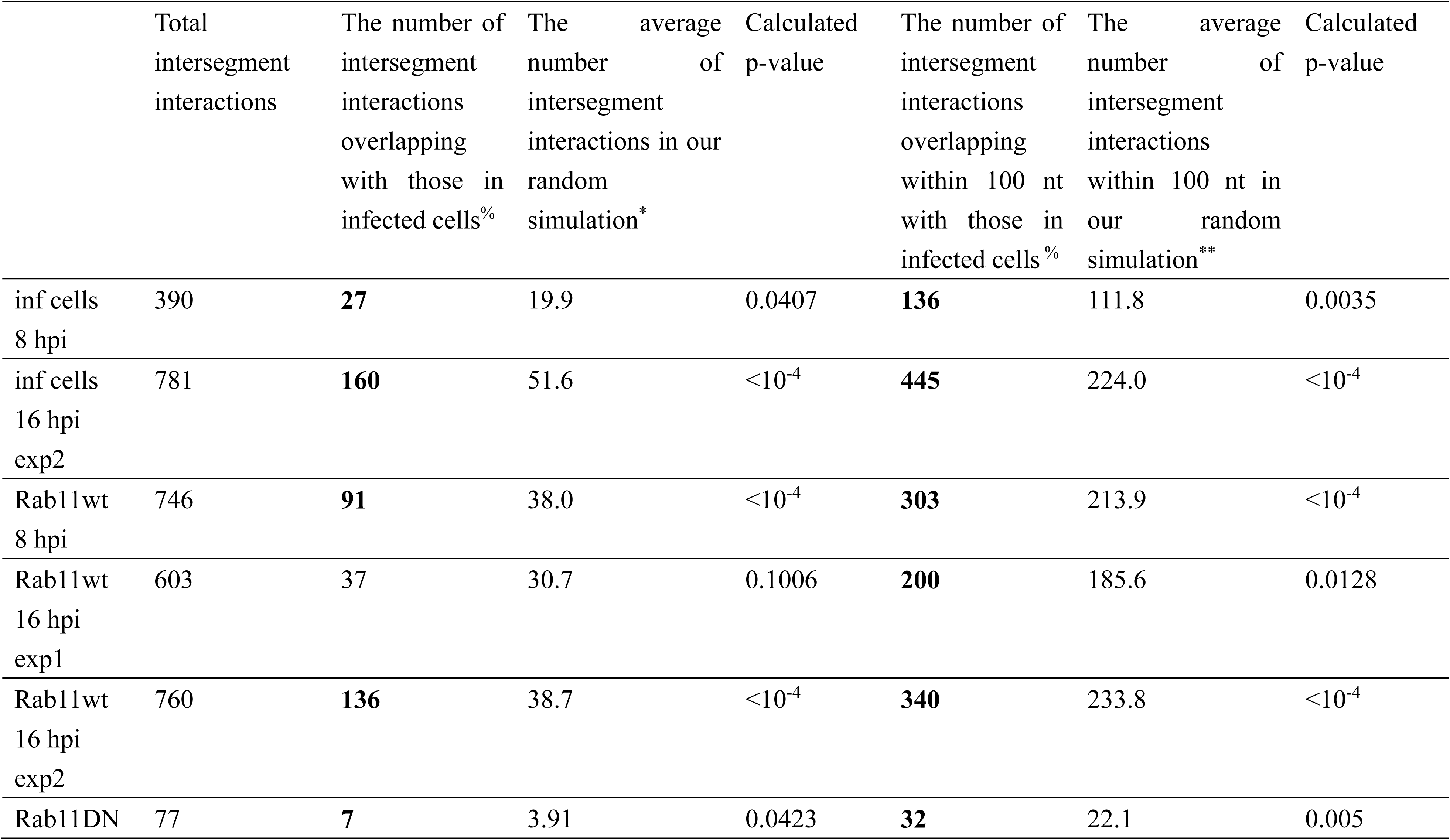
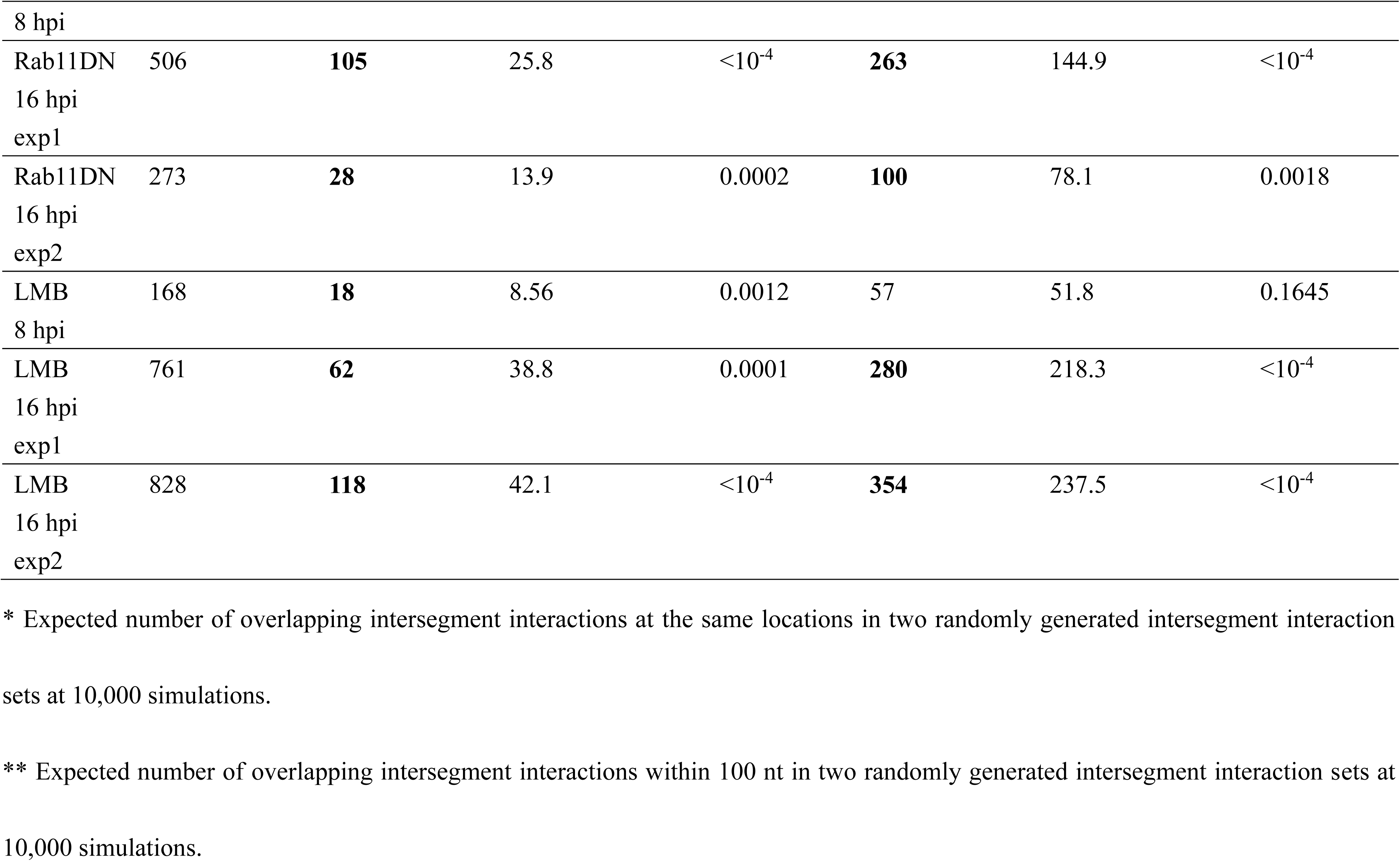
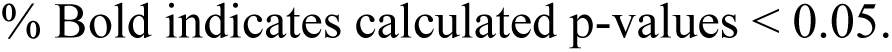
Comparison with intersegment interactions in MDCK cells (16 hpi, exp1)

Next, we examined whether the intersegment interactions in infected cells at 16 hpi are compatible with the prediction of RNA-RNA interactions. Among 431 intersegment interactions in infected cells at 16 hpi, 12 and 78 overlapped at the same locations and within 100 nt with predicted intersegment interactions, respectively. In our random simulation, average numbers of 7.8 and 60.4 intersegment interactions were reproduced at the same locations and within 100 nt, respectively, and the calculated p-value was less than 0.05 (*p* = 0.046 and 0.007, respectively) (Table 2). These results suggest that intersegment interactions in infected cells are partially formed by direct RNA interactions.

### Formation of viral inclusions in MDCK-Flag Rab11DN cells

It is hypothesized that intersegment interactions are formed in viral inclusions. Therefore, we aimed to identify intersegment interactions in viral inclusions. Before performing a comprehensive intersegment interaction analysis, we analyzed the formation of viral inclusions in MDCK-Flag Rab11 wild type (Rab11wt) and MDCK-Flag Rab11DN cells, since Rab11 is known to important for viral inclusion formation. MDCK-Flag Rab11wt and MDCK-Flag Rab11DN cells express Flag-tagged human Rab11A and Flag-tagged dominant negative (S25N) form of human Rab11A, respectively ^24^ (Figure S7A). Few infectious particles were detected in the supernatant at 8 hpi, while 10^4^ of infectious particles were detected in the supernatant from MDCK and MDCK-Flag Rab11wt cells at 16 hpi (Figure S7B). In contrast, the amount of infectious particles in the supernatant from MDCK-Flag Rab11DN cells at 16 hpi was approximately 1/10. These results suggest that viral budding increases after 8 hpi and confirm that Rab11DN acts as a dominant negative for virus propagation. Viral inclusion formation was examined by vRNA and Rab11 localization. Viral inclusions were observed in MDCK and MDCK-Flag Rab11wt cells, while those were not observed in MDCK-Flag Rab11DN cells at 8 hpi (Figure 3A and 3B). However, viral inclusions were observed in MDCK-Flag Rab11DN cells at 16 hpi, and neither endogenous Rab11 nor Flag-Rab11DN were colocalized at the viral inclusions (Figure 3C). The number of cells expressing Rab11DN in which viral inclusions were formed was comparable to those of MDCK cells and cells expressing Rab11wt. Viral inclusions were also observed in MDCK cells transiently expressing GFP-Rab11DN infected with PR8 and WSN strain (Figure S8). However, viral inclusions were not observed in A549 cells transiently expressing GFP-Rab11DN at 16 hpi (Figure S8). These results suggest that viral inclusion formation in cells expressing Rab11DN is cell-type specific. To characterize the viral inclusions in MDCK-Flag Rab11DN cells, the number of particles in the cells, the area of particles, and the circularity of particles were analyzed. The number of particles in MDCK, MDCK-Flag Rab11wt, and MDCK-Flag Rab11DN cells was comparable (Figure 3D). The area of particles in MDCK-Flag Rab11wt cells was slightly decreased compared to those in MDCK and MDCK-Flag Rab11DN cells (Figure 3E). The circularity of particles in MDCK-Flag Rab11DN cells was decreased compared to those in MDCK and MDCK-Flag Rab11wt cells (Figure 3F). These results suggest that the shape of viral inclusions in MDCK-Flag Rab11DN cells differs from typical droplets. To exclude the possibility that Rab11C (Rab25) induces the formation of viral inclusion in cells without functional Rab11, MDCK Rab11KO and Rab11/25 KO cells were infected with IAV. Viral inclusions were observed in both MDCK Rab11KO and Rab11/25 KO cells at 16 hpi, suggesting that Rab25 was not involved in the formation of viral inclusions without Rab11 in MDCK cells (Figure S9).

**Figure 3.**
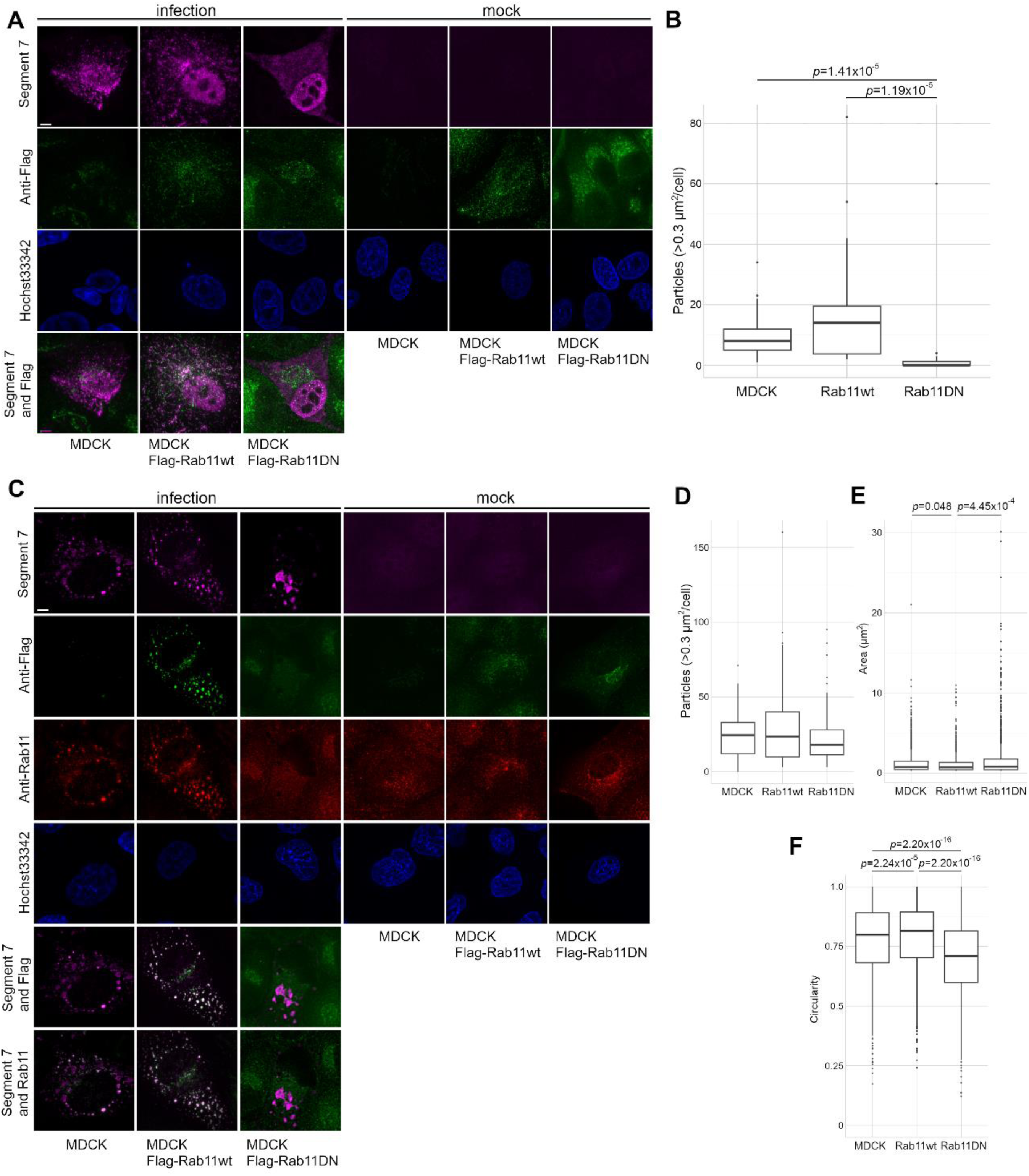
Viral inclusions in cells expressing Rab11 dominant negative at 16 hours post-infection. (A) Viral inclusion formation at 8 hpi. Cells were infected with IAV at an MOI of 1. Viral RNA (segment 7) and Flag-Rab11 in MDCK, MDCK-Flag Rab11 wild type (wt), and MDCK-Flag Rab11 dominant negative (DN) were visualized by in situ hybridization and immunofluorescence at 8 hpi. Bar: 5 µm. (B) Number of viral inclusions in infected cells. The number of viral RNA particles larger than 0.3 µm^2^ was measured. P-values were calculated using the Wilcoxon rank sum test and corrected using Bonferroni correction. The number of measured cells is 21 (MDCK) or 20 (Rab11wt and Rab11DN). (C) Viral inclusion formation at 16 hpi. Cells were infected with IAV at an MOI of 1. Viral RNA (segment 7), total Rab11 (endogenous and Flag-tagged), and Flag-Rab11 in MDCK, MDCK-Flag Rab11wt, and MDCK-Flag Rab11DN were visualized by in situ hybridization and immunofluorescence at 16 hpi. Rab11 was colored green in the merged images of segment 7 and Rab11. Bar: 5 µm. (D-F) Characteristics of viral inclusions in each cell. The number (D), area (E), and circularity (F) of viral RNA particles larger than 0.3 µm^2^ were measured. P-values were calculated using the Wilcoxon rank sum test and corrected using Bonferroni correction. Number of measured cells is 70 (MDCK), 44 (Rab11wt), or 58 (Rab11DN). Number of measured particles is 1729 (MDCK), 1378 (Rab11wt), or 1392 (Rab11DN).

To investigate whether viral inclusions in cells expressing Rab11DN have droplet properties, infected cells were treated with 1,6 hexanediol, and the vRNP particle size was examined. The number of particles in MDCK, MDCK-Flag Rab11wt, and MDCK-Flag Rab11DN cells was decreased by the treatment of 1,6 hexanediol (Figures 4A-D). Next, we analyzed the coalescence and splitting of viral inclusions in infected cells expressing Rab11DN. PA-GFP virus, which encodes the GFP-tagged polymerase subunit PA gene, was used for live cell imaging of vRNPs. Similar to infected cells expressing Rab11wt, coalescence and splitting of viral inclusions were observed in infected cells expressing Rab11DN (VideoS1, S2, and S3). These results indicate that vRNP can form viral inclusions without functional Rab11 in infected cells, although the timing of viral inclusion formation is delayed. To analyze the properties of viral inclusions in cells expressing Rab11DN in more detail, the dissolution of viral inclusions in cells expressing Rab11DN by hypotonic solution treatment was investigated. A previous report has shown that hypotonic solution treatment clearly reduces viral inclusions in infected cells more than 1,6-hexanediol treatment ^32^. Viral inclusions in MDCK cells and cells expressing Rab11wt were dissolved by hypotonic solution treatment, while the number of viral inclusions in cells expressing Rab11DN was not significantly changed by the treatment (Figures 4E-H). These results suggest that viral inclusions in cells expressing Rab11DN are resistant to hypotonic solutions treatment.

**Figure 4.**
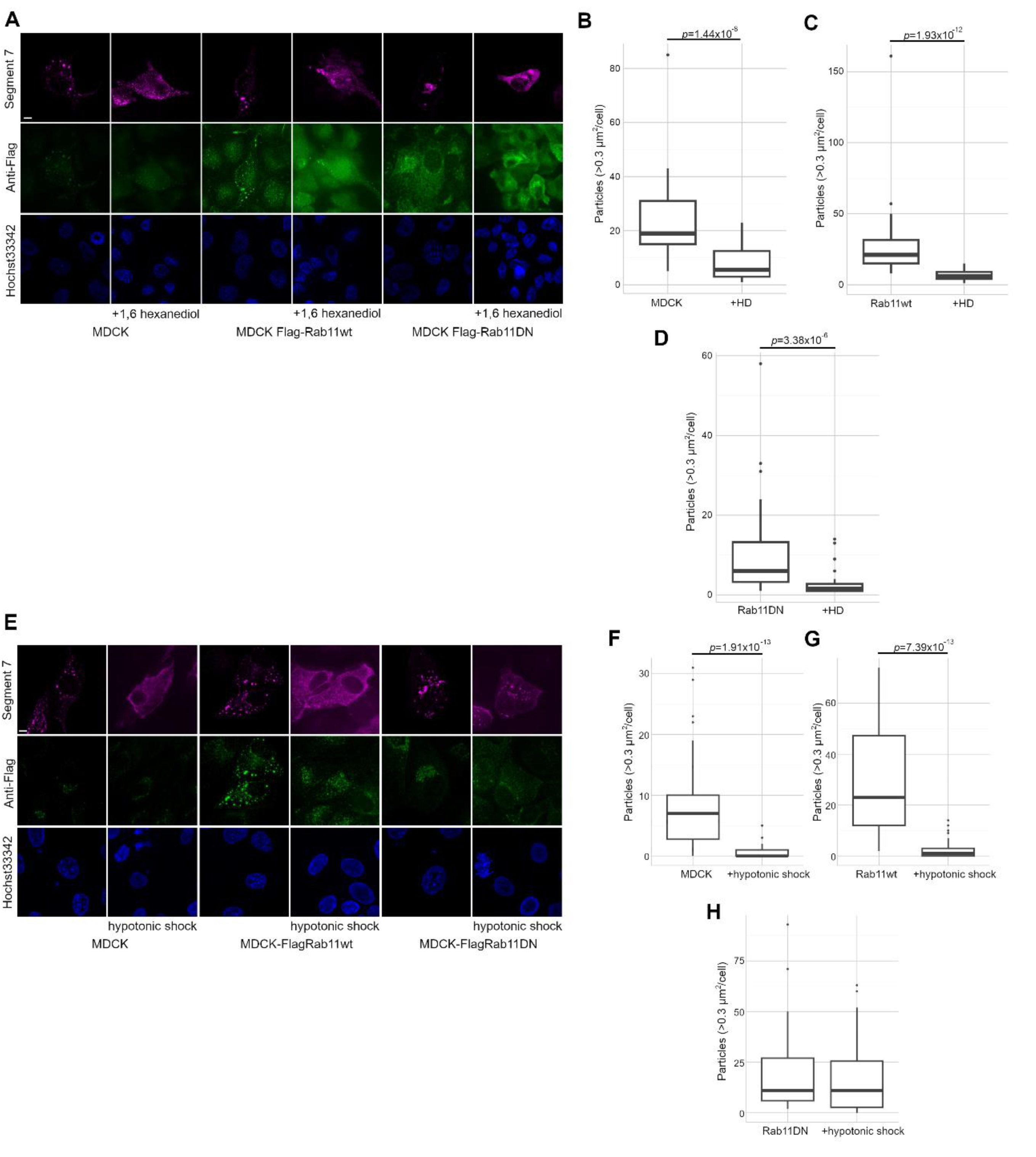
Dissolution and resistance of viral inclusions in cells expressing Rab11DN by 1,6-hexanediol and hypotonic solution. (A) Localization of vRNP and Rab11 in cells treated with 1,6-hexanediol. MDCK, MDCK-Flag Rab11wt, and MDCK-Flag Rab11DN were infected with IAV at an MOI of 1. Cells were treated with 4% 1,6-hexanediol for 5 min at 16 hpi. Viral RNA (segment 7) and Flag-Rab11 in MDCK, MDCK-Flag Rab11wt, and MDCK-Flag Rab11DN were visualized by in situ hybridization and immunofluorescence. Bar: 5 µm. (B-D) Particle size of viral inclusions in cells treated with 1,6-hexanediol (1,6 HD). The number of viral RNA particles larger than 0.3 µm^2^ were measured in MDCK (B), MDCK-Flag Rab11wt (C), and MDCK-Flag Rab11DN (D). P-values were calculated using the Wilcoxon rank sum test. Number of measured cells is 32 (MDCK), 32 (MDCK+1,6 HD), 31 (Rab11wt), 23 (Rab11wt+1,6 HD), 30 (Rab11DN), or 30 (Rab11DN+1,6 HD). (E) Viral inclusion in cells treated with hypotonic solution. Cells were infected with IAV at an MOI of 1. Viral RNA (segment 7) and Flag-Rab11 in MDCK, MDCK-Flag Rab11wt, and MDCK-Flag Rab11DN were visualized by in situ hybridization and immunofluorescence at 16 hpi with or without hypotonic solution treatment (37 °C for 5 min). Bar: 5 µm. (F-H) The particle size of viral inclusions in cells treated with hypotonic solution. The number of viral RNA particles larger than 0.3 µm^2^ was measured in MDCK (F), MDCK-Flag Rab11wt (G), and MDCK-Flag Rab11DN (H). P-values were calculated using the Wilcoxon rank sum test. The number of measured cells is 64 (MDCK), 47 (MDCK, hypotonic shock), 32 (Rab11wt), 55 (Rab11wt, hypotonic shock), 41 (Rab11DN), or 56 (Rab11DN, hypotonic shock).

Next, we determined the movement of viral inclusions and the exchange of vRNP from viral inclusions in MDCK-Flag Rab11DN cells. Movement of PA-GFP inclusions in MDCK, MDCK-Flag Rab11wt, and MDCK-Flag Rab11DN were tracked. The average movement speed of the PA-GFP inclusions in MDCK-Flag Rab11DN cells was decreased compared to that in MDCK and MDCK-Flag Rab11wt cells (Figures 5A and 5B). These results suggest that the movement of viral inclusions is restricted by the lack of functional Rab11 proteins in viral inclusions. To examine the exchange of vRNP from viral inclusions, we performed FRAP analysis using the PA-GFP virus. The rate and speed of PA-GFP fluorescence recovery of viral inclusions in MDCK, MDCK-Flag Rab11wt, and MDCK-Flag Rab11DN were comparable (Figure 5C). These results suggest that the exchange rate of vRNPs from viral inclusions without Rab11 is comparable to that with Rab11. We next examined the diffusion of vRNPs in viral inclusions with or without Rab11. PA-GFP in viral inclusions was partially photobleached, and fluorescence recovery of the region was observed (Figure 5D). Although no significant differences were observed due to the diversity of viral inclusions, there was a trend toward slower diffusion of PA-GFP in viral inclusions in cells expressing Rab11DN (Figure 5D and 5E). Importantly, in our system, viral inclusion formation is delayed by the lack of Rab11 but not impaired, as viral inclusions are observed in MDCK Rab11DN at 16 hpi without colocalizing with Rab11. The inclusions have different material properties in Rab11DN conditions, as these do not respond to hypotonic shock and are stiffer despite being able to fuse and divide, dynamically move, and responding to 1,6 hexanediol. Together, the findings suggest that inclusions in Rab11DN are stiffer than in Rab11WT conditions, still retaining a viscoelastic phenotype.

**Figure 5.**
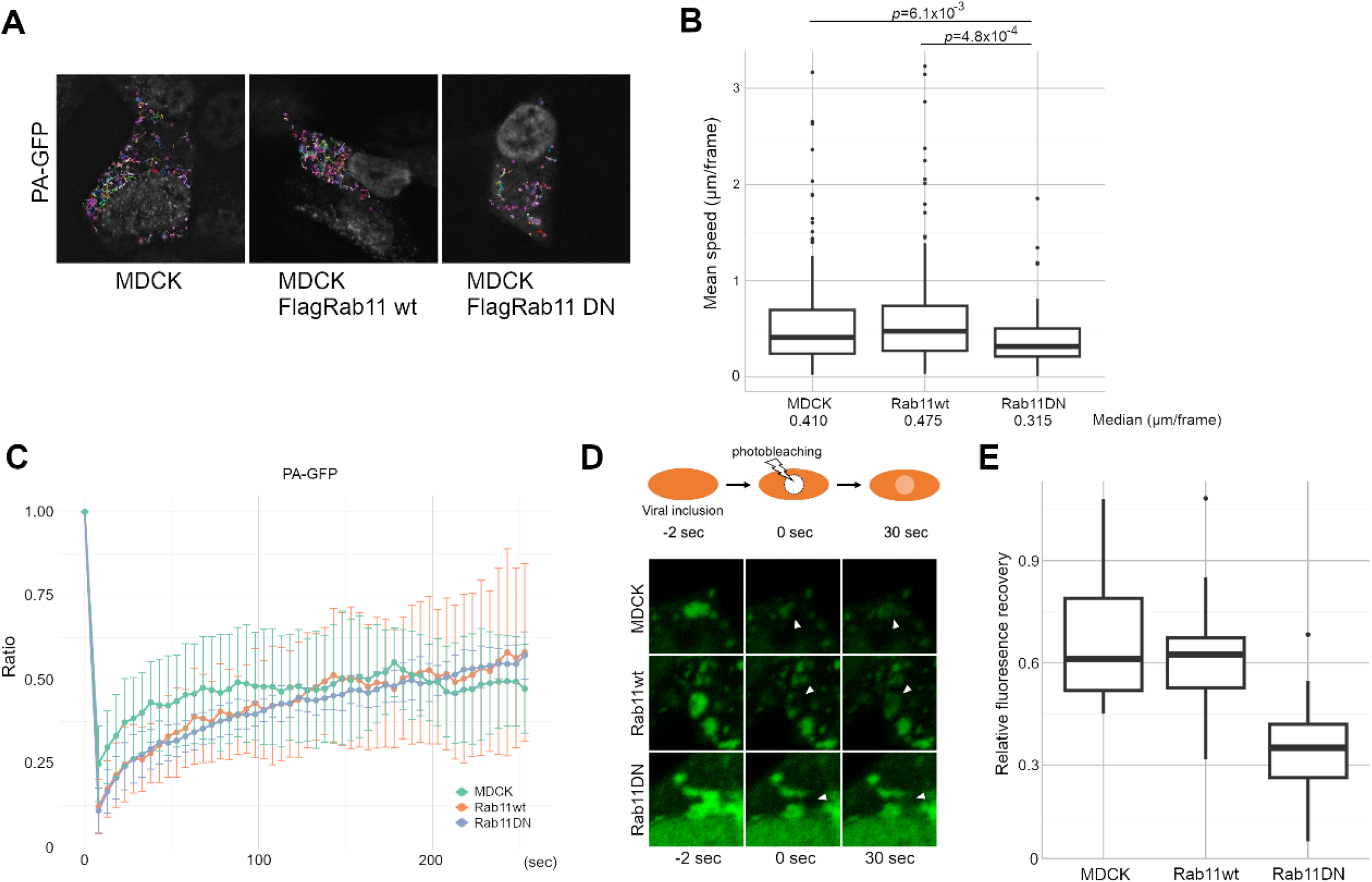
Properties of viral inclusions in cells expressing Rab11DN. (A) Tracking of viral inclusions. Viral inclusions in MDCK, MDCK-Flag Rab11wt, and MCK-Flag Rab11DN at 16 hpi were tracked by PA-GFP. (B) Mean speed of viral inclusions. The migration speed of viral inclusions was measured. P-values were calculated using the Wilcoxon rank sum test and corrected using Bonferroni correction. Number of measured particles is 203 (MDCK), 134 (Rab11wt), or 100 (Rab11DN). (C) FRAP assay of viral inclusion component proteins. FRAP assay of PA-GFP in viral inclusions was performed in MDCK, MDCK-Flag Rab11wt, and MDCK-Flag Rab11DN cells infected with PA-GFP virus at 16 hpi. The graph shows the average and standard deviation of fluorescence intensity of 5 droplets. (D) Diffusion of Rab11 in viral inclusions. An internal region of viral inclusion was photobleached, and the fluorescence recovery in viral inclusions was observed. MDCK, MDCK Flag-Rab11wt, and MDCK Flag-Rab11DN cells were infected with the PA-GFP virus, and cells at 16 hpi were treated with 10 µg/ml of nocodazole for 1 h to reduce the movement of viral inclusions. Viral inclusions before photobleaching (-2 sec), just after photobleaching (0 sec), and after 30 sec of photobleaching were shown. Arrowheads indicate a quenched area. (E) Fluorescence recovery rate after 30 sec of photobleaching. The relative fluorescence recovery in viral inclusions after 30 sec of photobleaching was calculated. Number of measured viral inclusions is 7 (MDCK), 7 (Rab11wt), or 9 (Rab11DN).

### Intersegment interaction in infected cells expressing Rab11 dominant negative

Viral inclusions are thought to be a site of segment assembly in infected cells, and we show that viral inclusions were formed in cells expressing Rab11DN even though the properties of viral inclusions without Rab11 were different from those with Rab11. Thus, to determine whether intersegment interactions occur in viral inclusions with or without Rab11, comprehensive intersegment interactions in cells expressing Rab11wt and Rab11DN were identified. MDCK-Flag Rab11wt and MDCK-Flag Rab11DN cells were infected with IAV, and the customized LIGR-seq was performed at 8 and 16 hpi. We identified 746 and 77 intersegment interactions in cells expressing Rab11wt and Rab11DN at 8 hpi, respectively (Figures 6A, 6B, S10A, and S11A). Among 746 intersegment interactions in cells expressing Rab11wt at 8 hpi, 75 and 291 were reproduced at the same locations and within 100 nt in the virion, respectively. In our random simulation, average numbers of 58.1 and 266.6 intersegment interactions were reproduced at the same locations and within 100 nt, respectively, and the calculated p-values were less than 0.05 (*p* = 0.009 and 0.026, respectively) (Table 1). Among 77 intersegment interactions in cells expressing Rab11DN at 8 hpi, 6 and 22 were reproduced at the same locations and within 100 nt in the virion, respectively. In our random simulation, average numbers of 5.95 and 27.5 intersegment interactions were reproduced at the same locations and within 100 nt, respectively, and the calculated p-values were more than 0.05 (*p* = 0.390 and 0.883, respectively) (Table 1). These results suggest that intersegment interactions in cells expressing Rab11wt at 8 hpi significantly overlap with those in the virion, but those in cells expressing Rab11DN do not. Moreover, we identified 603 and 506 intersegment interactions in cells expressing Rab11wt and Rab11DN at 16 hpi, respectively (Figures 6C, 6D, S10B, S10C, S11B, and S11C). Among 603 intersegment interactions in cells expressing Rab11wt at 16 hpi, 61 and 244 were reproduced at the same locations and within 100 nt in the virion, respectively. In our random simulation, average numbers of 46.9 and 215.6 intersegment interactions were reproduced at the same locations and within 100 nt, respectively. The calculated p-values were less than 0.05 (*p* = 0.015 and 0.005, respectively), suggesting that intersegment interactions in cells expressing Rab11wt at 16 hpi significantly overlap with those in the virion. Among 506 intersegment interactions in cells expressing Rab11DN at 16 hpi, 131 and 318 were reproduced at the same locations and within 100 nt in the virion, respectively. In our random simulation, average numbers of 39.4 and 180.9 intersegment interactions were reproduced at the same locations and within 100 nt, respectively, and the calculated p-values were less than 0.05 (*p* < 10^-4^) (Table 1). We performed two independent experiments on cells expressing Rab11wt and Rab11DN at 16 hpi, and similar results were obtained (Table 1). These results suggest that intersegment interactions in cells expressing Rab11DN at 16 hpi significantly overlap with those in the virion. Taken together, our comprehensive intersegment interaction analyses indicate that the formation of intersegment interactions is delayed in cells expressing Rab11DN and depends on viral inclusion formation with or without Rab11.

**Figure 6.**
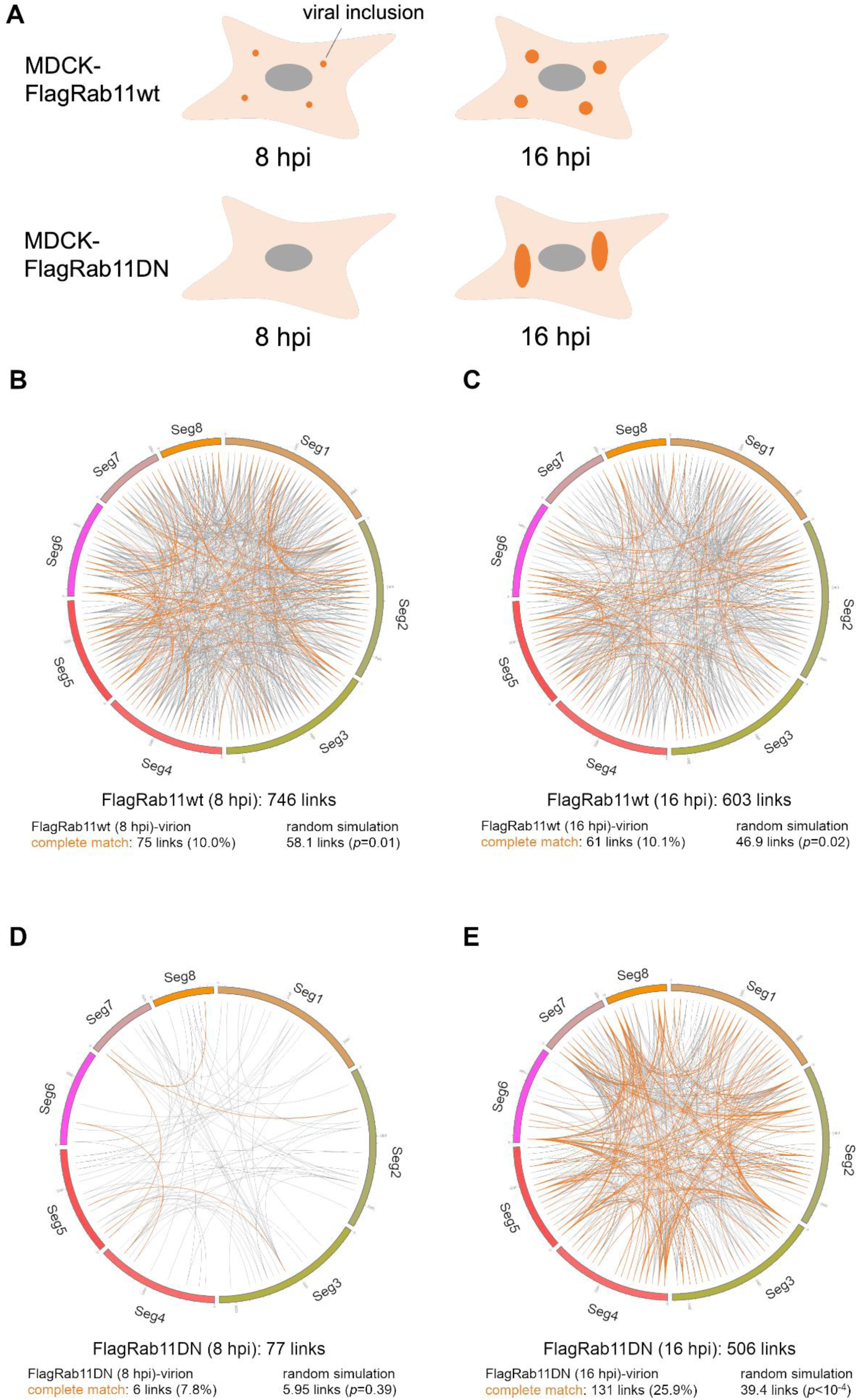
Intersegment interactions in infected cells expressing Rab11wt and Rab11DN. (A) Representative depiction of cells subjected to LIGR-seq. (B-D) Circos plots of the intersegment interactions in MDCK-Flag Rab11wt cells at 8 hpi (B) and 16 hpi (C) and MDCK-Flag Rab11DN cells at 8 hpi (D) and 16 hpi (E). Intersegment interactions identified in both the virion and the cell are indicated by orange lines. “Random simulation” means the number of overlapping intersegment interactions between two randomly generated intersegment interaction sets.

### Intersegment interactions under artificially increased local vRNP concentration

Based on our results, we hypothesize that an increase in local vRNP concentrations induces the formation of intersegment interaction networks. To test the hypothesis that intersegment interaction occurs in a vRNP concentration-dependent manner, we artificially increased vRNP concentration in the nucleus where segments are not colocalized under normal conditions ^25–27^. We identified comprehensive intersegment interactions in infected cells treated with leptomycin B (LMB), which inhibits the nuclear export of vRNP. Viral RNAs were accumulated in the nucleus in cells treated with LMB (Figure 7A). Viral segments were uniformly distributed in the nucleus by the treatment with LMB to 8 hpi and were not co-concentrated, while those were co-concentrated in some areas of the nucleus by the treatment with LMB to 16 hpi (Figure 7B). Next, the customized LIGR-seq was performed on infected cells with LMB to 8 and 16 hpi. We identified 168 intersegment interactions in infected cells with LMB to 8 hpi, and 9 and 64 intersegment interactions were reproduced at the same locations and within 100 nt in the virion, respectively (Figures 7C and S12A). In our random simulation, average numbers of 13.1 and 60.0 intersegment interactions were reproduced at the same locations and within 100 nt, respectively, and the calculated p-values were more than 0.05 (*p* = 0.852 and 0.242, respectively) (Table 1). The intersegment interactions in infected cells with LMB to 8 hpi significantly overlapped with those in infected MDCK cells at 16 hpi (*p* = 0.001) (Table 3), despite not significantly overlapping with those in the virion. This result suggests that intersegment interactions with roles other than segment assembly are formed in infected cells. We identified 761 intersegment interactions in infected cells with LMB to 16 hpi, and 136 and 394 intersegment interactions were reproduced at the same locations and within 100 nt in the virion, respectively (Figures 7D and S12B). In our random simulation, average numbers of 59.2 and 272.0 intersegment interactions were reproduced at the same locations and within 100 nt, respectively, and the calculated p-values were less than 0.05 (*p* < 10^-4^) (Table 1). We performed two independent experiments on LMB-treated cells at 16 hpi, and similar results were obtained (Table 1). These results suggest that intersegment interactions overlapped with those in the virion are formed under artificially increased local vRNP concentration.

**Figure 7.**
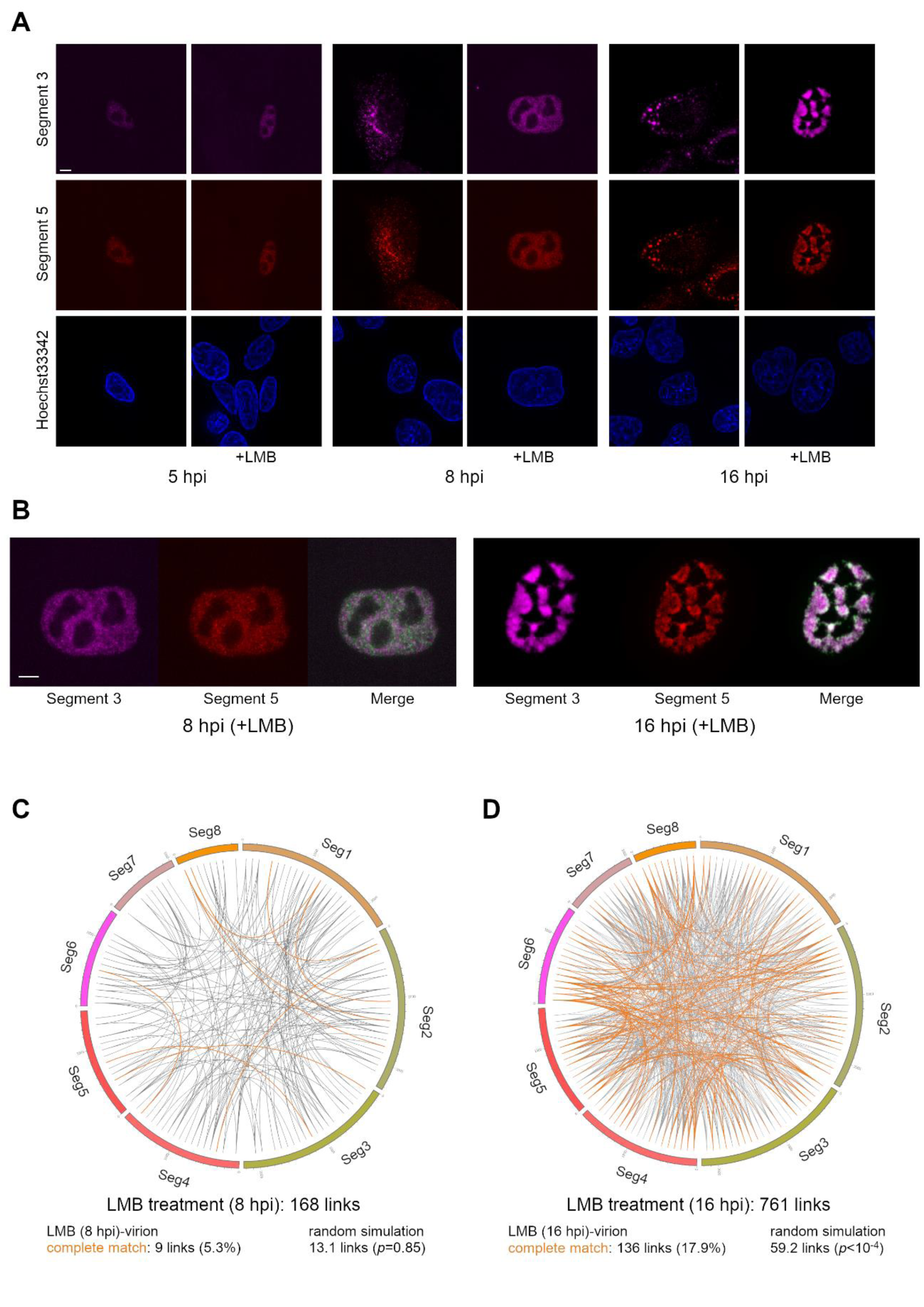
Intersegment interactions with artificially increased local concentrations of vRNA. (A) Localization of vRNAs in cells treated with leptomycin B. MDCK cells were infected with IAV at an MOI of 1 and treated with or without 25 ng/ml leptomycin B (LMB). Segments 3 and 5 were visualized by in situ hybridization at 5, 8, and 16 hpi. Bar: 5 µm. (B) Colocalization of segments in LMB-treated cell nuclei at 16 hpi. Enlarged views of segments 3 and 5 in LMB-treated cell nuclei at 8 and 16 hpi and merged images were shown. Segment 5 is colored green and white areas indicate colocalization of segment 3 and 5 in the merged images. Bar: 5 µm. (C) Circos plots of the intersegment interactions in cells treated with LMB at 8 (C) and 16 hpi (D). Intersegment interactions identified in both the virion and the cell are indicated by orange lines. “Random simulation” means the number of overlapping intersegment interactions between two randomly generated intersegment interaction sets.

## Discussion

It has been hypothesized that the eight IAV genome segments are selectively bundled by intersegment RNA-RNA interaction networks, and that viral inclusions serve as cellular compartments where IAV segments are bundled in infected cells. To investigate this, we analyzed comprehensive intersegment interactions in infected cells with or without functional Rab11 by customized LIGR-seq. This study focused on the overall intersegment interaction network, rather than individual interactions, and examined overlaps with intersegment interactions in the virion, since the intersegment interaction network is redundant and complex ^17–19^. Our findings show that intersegment interactions overlapping with those in the virion are formed in infected cells where viral inclusions are formed and in the nuclei of infected cells treated with LMB. These results suggest that the intersegment interaction networks of the IAV genome are formed at viral inclusions due to an increase in the local concentration of vRNP.

We identified intersegment interactions in the virion and infected cells by the customized LIGR-seq. By combining statistical analyses, we have enhanced the reliability of our data. We compared our LIGR-seq data for the virion with the SPLASH data of the PR8 strain ^17^, since intersegment interactions vary between strains. The sequences of the PR8 strain we used and those used in the SPLASH experiment are 99.14% identical. The intersegment interactions identified by LIGR-seq overlapped with those identified by SPLASH (Figure S4). Moreover, the intersegment interactions identified by LIGR-seq and SPLASH overlapped with predicted intersegment interactions (Table 2). These findings confirm the reliability of our customized LIGR-seq data. The purified virion used in our LIGR-seq was amplified in the allantoic sacs of chick embryos, whereas that used in SPLASH was amplified in MDCK cells. Intersegment interactions are expected to be identical in different cell types if the viral strains are the same.

In previous studies, viral inclusions were not detected in cells expressing Rab11DN ^22–24,30,32,37,38^. However, our results showed that viral inclusions were formed in MDCK cells expressing Rab11DN at 16 hpi (Figure 3C). Viral inclusions were formed in Rab11KO (both Rab11A and Rab11B KO) and Rab11/25 KO cells (Figure S9), suggesting that other Rab11 family proteins do not rescue viral inclusion formation in cells expressing Rab11DN. Viral protein expression in cells expressing Rab11DN or Rab11 knock-down cells was comparable to that in normal cells ^24,37^, and viral inclusions were not detected in cells expressing Rab11DN at 8 hpi while those were formed in cells expressing Rab11wt (Figure 3A). Our results indicate that functional Rab11 could facilitate the formation of viral inclusions. The delay in viral inclusion formation is one of the factors contributing to the delay of the viral propagation in MDCK Rab11DN cells at 8 hpi. Notably, viral inclusions were not observed in A549 cells expressing Rab11DN (Figure S8). This result indicates that viral inclusion formation in cells lacking functional Rab11 is cell-type specific. The cellular state would be changed at the late stage of infection in MDCK cells lacking functional Rab11, and vRNPs bind to other host factors that induce droplet formation of vRNP. In fact, viral inclusion in cells expressing Rab11DN had droplet characteristics, but the properties were slightly different compared to normal viral inclusion (Figures 3F, 4B, and 5B). Rab11 binds to actin and microtubule motor proteins directly and via FIPs ^39^, and IAV infection alters the binding of Rab11 to these motor proteins ^40^. Viral inclusions without Rab11 could not bind to motor proteins, leading to differences in the movement and circularity of viral inclusions compared to those with Rab11. Viral propagation in cells expressing Rab11DN was reduced compared to wild type cells, despite the formation of viral inclusions in MDCK Rab11DN cells (Figure S7B). This suggests that one of the functions of Rab11 in viral inclusions is to facilitate their transport. A previous report indicates that ATG9A impacts the affinity of viral inclusions to microtubules and that the shape of viral inclusions is incorrect in ATG9A knock-down cells ^41^. This phenotype is identical to viral inclusions without Rab11. Furthermore, the lack of Rab11 or ATG9A affects viral genome bundling ^38,41^. Transport of viral inclusions by microtubules may be necessary for maturation of viral inclusions and correct assembly of viral genome in viral inclusions.

The intersegment interactions were redundant, and a single region coordinates multiple interactions with other segments (hotspots). These redundant intersegment interactions and hotspots can be explained by varying interactions in each virion. In our analysis, hotspots were found at the 3′ and 5′ ends and in the middle of the segment both in virions and infected cells (Figure S3). These hotspot distributions were not observed in the interaction prediction (Figure S3). These findings suggest that intersegment interactions are topologically constrained. There are two possible topological constraints: vRNP formation and vRNP orientation. In the vRNP state, the base of the vRNA is exposed externally, allowing RNA-RNA interactions to form. However, the structure of vRNPs prevents the formation of long RNA-RNA interactions between segments. To incorporate this constraint into predicting intersegment interactions, we used an algorithm that predicts short RNA-RNA interactions. vRNPs are primarily packaged in a head-to-tail, and to some extent, in a tail-to-head orientation ^35^. The hotspot distribution in infected cells is comparable with that in the virion, indicating that vRNP orientation is at least partially aligned in infected cells. In infected cells, vRNP orientation alignment likely occurs prior to the formation of intersegment interactions.

Intersegment interactions in cells treated with LMB and in cells expressing Rab11DN at 8 hpi overlapped with those in infected cells at 16 hpi (Table 3). However, vRNPs were not concentrated in cells treated with LMB or cells expressing Rab11DN at 8 hpi, and intersegment interactions in those cells did not overlap with those in the virion. These findings suggest that certain intersegment interactions may have functions other than viral genome packaging into the virion. In addition, intersegment interactions in infected cells overlapped with predicted interactions, but the percentage of concordance was low compared to those between virion and predicted interactions (Table 2). In infected cells, it is likely that vRNPs are in close proximity to each other due to factors other than direct RNA-RNA interactions. These indirect RNA-RNA intersegment interactions would be resolved before budding, and the orientation of vRNPs is thought to align. Viral inclusion contains more than one set of eight segments, since their size is larger than that of a virion. The next question to be addressed is whether one set of eight segments is released from viral inclusions or whether viral inclusion collapses after transporting vRNPs to the budding site.

In conclusion, our data indicates that intersegment interaction networks are formed in viral inclusions. Enrichment of progeny vRNPs in viral inclusions is efficient for the formation of intersegment interaction networks. It is possible that the plasticity of the droplet is used to release one set of eight segments to the budding site, but this hypothesis needs to be analyzed.

## Materials and methods

### Cells, plasmids, and viruses

MDCK cells were obtained from the ATCC (CCL-34). MDCK-Flag Rab11wt and MDCK-Flag Rab11DN cells were provided by Dr. Momose ^24^. MDCK II, MDCK Rab11KO, and MDCK Rab11/25KO cells were obtained from the RIKEN BRC (RCB5148, RCB5112, and RCB5143) ^42^. Cells were maintained in a Dulbecco’s modified Eagle’s medium (DMEM) (Wako Pure Chemical Industries, Osaka, Japan) with high glucose concentration containing 10% fetal bovine serum and penicillin/streptomycin (Nacalai Tesque, Kyoto, Japan). HEK293T and A549 cells were provided by Dr. Kawaoka. Cells were maintained in a DMEM with high glucose concentration (Wako Pure Chemical Industries) containing 10% fetal bovine serum and penicillin/streptomycin.

The GFP-tagged human Rab11A expression vector (pCANeoAcGFP-hRab11A) was provided by Dr. Momose. To construct the GFP-Rab11DN expression vector, an inverted PCR was performed using the pCANeoAcGFP-hRab11A vector as a template and KOD One (Toyobo, Osaka, Japan) with specific primer sets. After DpnI treatment, phosphorylation, ligation and transformation into an *Escherichia coli* Mach 1 was performed. Transient transfection to MDCK and A549 cells was performed using Lipofectamine 3000 (Thermo Fisher Scientific, Waltham, MA). After 30 h transfection, cells were infected with IAV.

Influenza virus A/PR/8/34 (H1N1) (PR8) and A/WSN/1933 (H1N1) (WSN) were prepared using a reverse genetics approach ^43,44^ and grown in MDCK cells. For preparing purified virion, PR8 was grown in the allantoic sacs of 11-day-old chick embryos at 35.5°C for 48 h. The purified virion was prepared as previously described ^45^. PA-GFP virus was prepared using a reverse genetics approach as previously described ^32^. Reverse genetics plasmids were provided by Dr. Fouchier ^46^ and Dr. Amorim ^32^. PA-GFP virus was rescued in HEK293T cells and amplified in MDCK cells.

### In situ hybridization and immunofluorescence

In the 1,6-hexanediol or hypotonic solution treatment experiment, infected cells at 16 hpi were treated with 4% 1,6-hexanediol (Tokyo Chemical Industry, Tokyo, Japan) in MEM or 1/10 diluted MEM for 5 min at 37 °C. Cells were fixed with 4% paraformaldehyde in PBS(-) at room temperature for 10 min, followed by permeabilization by 0.2% NP-40/PBS(-) at room temperature for 10 min. The cells were immersed in 0.2% BSA/PBS(-) at room temperature for 1 h and incubated with appropriate primary antibody at room temperature for 1 h. The cells were washed with PBS and incubated with appropriate secondary antibody conjugated with Alexa dye at room temperature for 1 h. The cells were washed with PBS(-) and re-fixed with 4% paraformaldehyde in PBS(-) at room temperature for 10 min. The cells were immersed in 2xSSC-10% formamide at 37°C for 10 min and incubated with hybridization solution (2xSSC, 10% formamide, 10% dextran sulfate, 0.02% BSA, 10 µg tRNA, and 160 nM FISH probe) at 37°C for 16 h. The cells were washed with 2xSSC-10% formamide at 37°C for 30 min, and the cell nuclei were stained with Hoechst33342 diluted with 2xSSC at room temperature for 10 min. The cells were observed under fluorescence microscopy. Mouse monoclonal anti-Flag (Sigma-Aldrich, ST. Louis, MO) and rabbit polyclonal anti-Rab11A (Proteintech, Rosemont, IL) antibodies were used for primary antibodies. Anti-mouse IgG Alexa 488 and anti-rabbit IgG Alexa 546 (Thermo Fisher Scientific) were used for secondary antibodies. Stellaris RNA FISH probes for segment 5 vRNA labeled with CAL Fluor Red 610 and segment 3 and 7 vRNAs labeled with Quasar 670 were purchased from LGC Biosearch Technologies (Hoddesdon, UK). Images were acquired with a THUNDER 3D Live Cell system (Leica Microsystems, Wetzlar, Germany). Z-stack scans were performed at 0.2 µm intervals, and images of maximum intensity projections and THUNDER computational clearing are shown. The number, area, and circularity of particles in the infected cells were analyzed using ImageJ ^47^. Images were acquired using tile scan mode without Z-stack, and binarization thresholding was performed by MaxEntropy in ImageJ. Particles larger than 0.3 µm^2^ were analyzed as viral inclusion in accordance with the previous paper ^32^.

### Live cell imaging and FRAP

In FRAP experiments, images were acquired with A1R HD25 confocal microscopy system (Nikon, Tokyo, Japan). MDCK, MDCK-Flag Rab11wt, and MDCK-Flag Rab11DN cells were infected with PA-GFP virus, and the infected cells at 16 hpi were photobleached and imaged for 5 min at 5-second intervals under 37°C and 5% CO_2_ conditions. In fluorescence recovery of a viral inclusion internal region observation, infected cells were photobleached and imaged for 1 min at no interval. Intensities of the photobleached region and unbleached internal region were measured by NIS-Elements AR (Nikon) and relative fluorescence recovery was calculated from these intensities. In live cell imaging experiments, images were acquired and deconvolved with a THUNDER 3D Live Cell system. MDCK cells were infected with the PA-GFP virus, and the infected cells at 16 hpi were imaged with a Z-stack scan (4 slices) for 3 min at 5-second intervals under 37°C and 5% CO_2_ conditions. Images of maximum intensity projections and THUNDER computational clearing were used for particle tracking. Particle tracking was performed using the trackMate plugin ^48^ in Fiji ^49^. The tracking algorithm used the simple LAP tracker.

### Customized LIGR-seq

For preparing cross-linked vRNA from virion, the purified virion in PBS(-) was treated with 100 µg/ml AMT (Sigma-Aldrich) and cross-linked using 365 nm UV for 20 min (1.56 J/cm^2^). After AMT treatment, the virion was treated with Proteinase K in PBS-0.1% SDS at 37 °C for 30 min. The viral RNA was extracted with phenol/chloroform. For preparing cross-linked vRNA from infected cells, cells (5×10^6^ cells in 100 mm dish) were infected with IAV at an MOI of 3. The infected cells were washed with PBS(-) and treated with 100 µg/ml AMT in PBS(-) using 365 nm UV for 30 min (2.34 J/cm^2^). The AMT solution was changed to a new one at 15 min after UV irradiation. Total RNA was extracted using the ISOGEN reagent (NIPPON GENE, Tokyo, Japan). The cross-linked vRNA was digested with the NEBNext Magnesium RNA Fragmentation Module (New England Biolabs, Ipswich, MA) at 94°C for 2.5 min. The fragmented RNA was treated with 15 U of CIAP (Takara Bio, Otsu, Japan) at 37°C for 30 min, and phosphorylation was carried out by the 10 U of T4 polynucleotide kinase (Toyobo) at 37°C for 1 h.

RNA pull-down procedure was performed using a modification of an RNA antisense purification method previously described for long noncoding RNA and influenza virus mRNA ^50,51^. cDNA of the PR8 strain was synthesized with the Uni12 primer using ReverTra Ace (Toyobo). The fragments of each segment were amplified using Taq polymerase (Roche, Basel, Switzerland) and specific primers with a linker sequence designed to amplify every 120 bp of the coding region with a 15-bp overlapping region (PCR primers were listed in the supplemental Table). To synthesize the biotinylated cDNA probe, a second PCR procedure was performed with the biotinylated linker primer using Taq polymerase. The biotinylated cDNA probe for each segment was mixed with the Dynabeads MyOne Streptavidin C1 (Thermo Fisher Scientific) and incubated on a rotating wheel at 37°C for 1 h in a LiCl hybridization buffer (10 mM Tris-HCl [pH 7.9], 500 mM LiCl, 1 mM EDTA, and 0.1 % NP-40). The beads were washed with the LiCl hybridization buffer and 1x SSPE and were suspended and incubated in 0.15 M NaOH for 10 min. After incubation, the mixture was neutralized with 100 mM Tris-HCl (pH 7.9) and 1.25 M AcOH. The beads were washed with 0.1 M NaOH and were suspended in the LiCl hybridization buffer. The cross-linked RNA was boiled at 85°C for 3 min and was added to the beads. The beads were incubated at 55°C in a Thermomixer (Eppendorf, Hamburg, Germany) at 1,500 rpm for 2 h. The beads were washed with 1x and 0.1x SSPE at 55°C. Proximity ligation was performed with 40 U of the T4 RNA ligase I (Takara Bio) at 16°C for 16 h in an RNA ligase buffer, mixing in a Thermomixer at 1,000 rpm for 15 sec every 15 min. The vRNA was eluted with 5 U of DNase I (Takara Bio) at 37°C for 30 min in a DNase buffer (40 mM Tris-HCl [pH 7.5], 8 mM MgCl_2_, and 5 mM DTT) and was extracted with phenol/chloroform. The eluted vRNA was treated with 10 U of RNaseR (Lucigen, Middleton, WI) in an RNaseR buffer (20 mM Tris-HCl [pH 8.0], 100 mM KCl, and 0.1 mM MgCl_2_) at 37°C for 30 min and was extracted with phenol/chloroform. The cDNA was synthesized with a random hexamer using the ReverTra Ace and the NEBNext mRNA Second Strand Synthesis Module (New England Biolabs). Sequencing libraries were constructed using the NEBNext UltraII DNA Library Prep Kit for Illumina (New England Biolabs). Sequencing was performed using a HiSeq2500 sequencer (Illumina, San Diego, CA) (2 × 100-bp PE) and a NovaSeq6000 sequencer (Illumina) (2 × 150-bp PE) and approximately 30 M reads were obtained per sample.

### Data analysis for segment interactions

Raw reads were cleaned and trimmed into the first 25 bases with Trimmomatic v0.36 ^52^. The cleaned reads were aligned to the PR8 genome using bowtie2 with default parameters^53^. Obtained sequencing reads were classified into two categories: intersegment and intrasegment interaction. We detected the former interaction by the paired-end reads that were mapped at two different segments and the latter ones that were mapped to the same segment at an inverted direction and a long insert length (more than 500 nt). The start positions of the selected pair-reads were counted in every 100 nt, and the contact map was constructed for the intra- and intersegment interactions. To normalize the biases, we adapted the iterative method which has been employed for the Hi-C analysis ^54^. The raw contact of each bin (*A_ij_*) in the contact map was divided by the sum of the contacts in the whole row (Σ_i_*A_ij_*) and the sum of the contacts in the whole column (Σ_j_*A_ij_*) to maintain matrix symmetry.

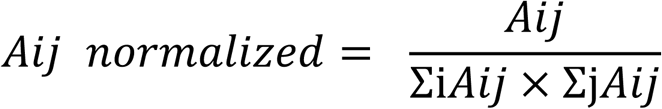

This calculation was repeated 50 times and convergence was confirmed. The normalized count in each 100 nt bin was referred to as the contact score. To discriminate accurately between the true and false signals, we performed LIGR-seq in the duplicate experiments and utilized an irreproducible discovery rate (IDR) ^55^. IDR compares a pair of ranked lists by contact scores and assigns IDR scores that reflect its reproducibility. The contact scores of all regions containing both intrasegment and intersegment regions of duplicate experiments were analyzed by IDR, and interactions with < 0.05 IDR (infected cell experiments) or < 0.01 IDR (virion experiments) were extracted. To exclude false positive interactions such as segment recombination during replication, we performed LIGR-seq experiments with or without RNA ligase treatment. Since a large part of non-crosslinked RNAs is digested by RNaseR, samples without RNA ligase treatment were used as negative controls instead of non-crosslinked samples. Any interaction bin detected in the samples without RNA ligase treatment was considered background signals and abandoned in the subsequent analyses. The intersegment interactions were visualized by Circos plot using Circos software ^56^. The intersegment interactions were predicted by RIblast (version 1.1) with default parameters ^36^.

To statistically evaluate the overlap of two experimentally obtained intersegment interactions, the expected overlapping intersegment interactions were simulated by randomly generating two sets of intersegment interactions, and the average number of overlapping intersegment interactions was calculated from 10,000 trials. The p-value was calculated with reference to the permutation p-value. The number of trials in the random simulation where the number of overlapping intersegment interactions exceeded the number of those identified by experiments was counted. The p-value was calculated by dividing the counted number of trials by the total number of trials (10,000). When the calculated p-value was less than 0.05, the intersegment interactions between the two conditions were considered overlapping.

### Western blotting

Cells were suspended in cell lysis buffer (20 mM Tris-HCl [pH 7.9], 100 mM NaCl, 1 mM EDTA, and 0.2% SDS) and sonicated. Cell lysates were separated by SDS-PAGE and detected by western blotting using an ImageQuant LAS 4000 (GE Healthcare, Milwaukee, WI). Rab11, Rab25, and tubulin were detected using rabbit polyclonal anti-Rab11A/B (proteintech), rabbit polyclonal anti-Rab25 (proteintech), and mouse monoclonal anti-α-tubulin (Sigma-Aldrich) antibodies. For fluorescence western blotting, cell lysates were separated by SDS-PAGE and detected using an Odyssey M (LI-COR Biosciences, Lincoln, NE). Rab11 and Flag-Rab11 were detected using rabbit polyclonal anti-Rab11A (proteintech) and rabbit polyclonal anti-Flag (MBL, Tokyo, Japan) antibodies. The total protein on the membrane was stained by Revert 520 Total Protein Stain Solution (LI-COR Biosciences). The band intensities were calculated using Empiria Studio (LI-COR Biosciences).

### Virus propagation

MDCK, MDCK-Flag Rab11wt, and MDCK-Flag Rab11DN were infected with PR8 strain at an MOI of 1 for 1 h at 37 °C in MEM(-). Cells were washed with MEM(-) and were incubated in MEM(-) pH 4.5 at 25 °C for 5 min to inactivate the remaining virus particles. Infected cells were suspended in MEM(-) and were incubated at 37 °C. The supernatant was collected at 2, 8, and 16 hpi and was treated with trypsin (final 0.66 µg/ml) at 37 °C for 30 min. The virus titer was determined by plaque assay.

## Acknowledgments

We thank Dr. Yoshihiro Kawaoka, Dr. Fumitaka Momose, Dr. Ronaldus Adrianus Maria Fouchier, and Dr. Maria João Amorim for kindly providing materials. We thank Ms. Yukiko Iwata for her technical support of experiments and Dr. Maria João Amorim for discussions and comments. This work was partly performed in the Cooperative Research Project Program of the Medical Institute of Bioregulation, Kyushu University. This work was supported by JSPS KAKENHI Grant Number 25871077, 15K21607, and 19K07598 to N.T., JSPS KAKENHI Grant Number 221S0002 and 16H06279 (PAGS), Japan Program for Infectious Diseases Research and Infrastructure from AMED Grant Number JP20wm0325008 to N.T., Takeda Science Foundation, GSK Japan Research Grant 2016, and the Waksman Foundation of Japan to N.T..

## Author contributions

Conceptualization, N.T.; Methodology, N.T., K.T., and K.K.; Formal Analysis, N.T. and R.K.K.; Investigation, N.T. and Y.G.; Writing – Original Draft, N.T.; Writing – Review & Editing, N.T. and R.K.K., Supervision, T.H. and K.K.; Founding Acquisition, N.T.

## Data availability

The sequence data have been deposited in DDBJ Sequence Read Archive. The BioProject number is PRJDB17853 and the DRR run numbers are DRR543786-DRR543841. Intersegment interaction data were provided in the supplemental data.

## Code availability

Source codes for matrix balancing normalization and random simulation are available on github (https://github.com/naoki-takizawa/flu_LIGR_seq).

---

**Figure S1.**
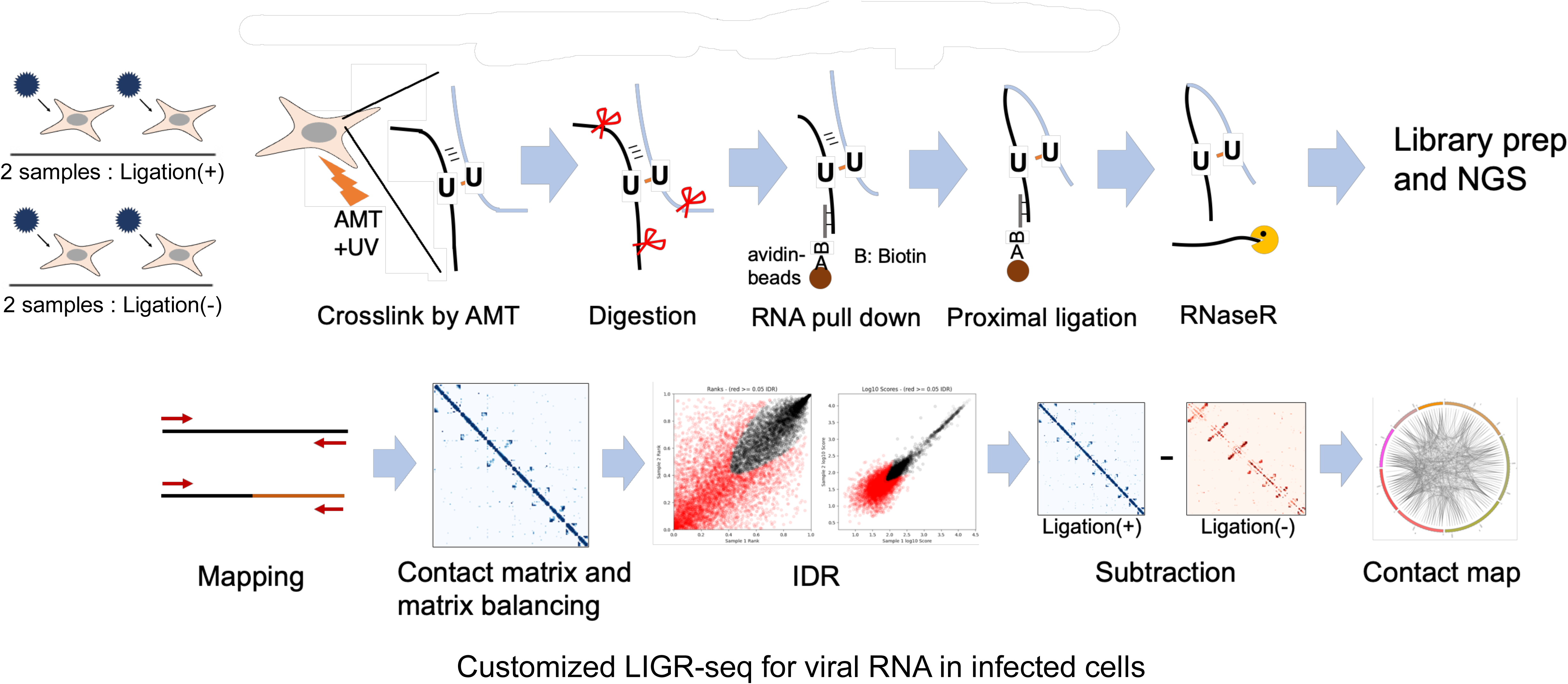
Schematic representation of the customized LIGR-seq and data analysis.

**Figure S2.**
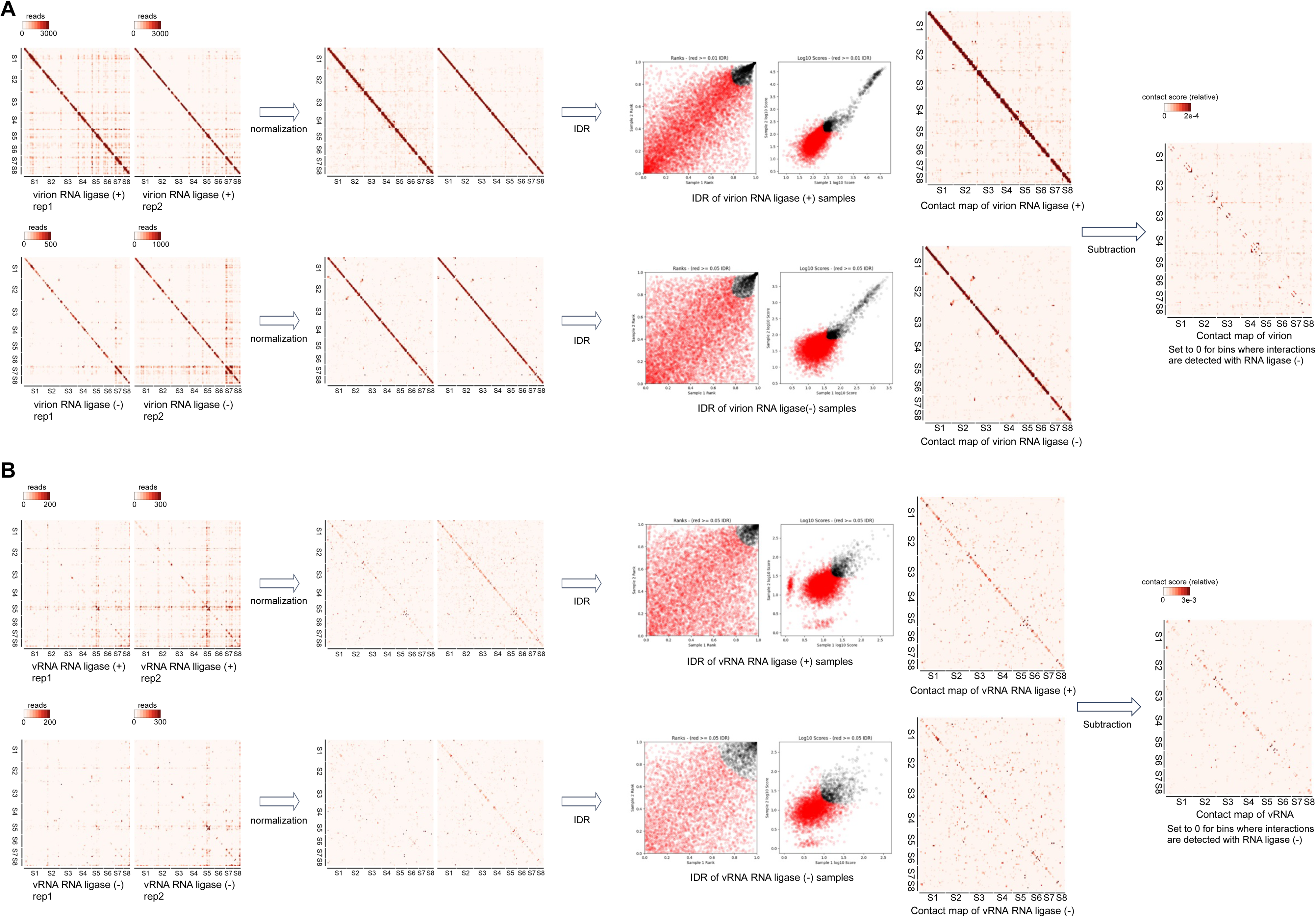
IDR outputs and contact maps of segment interactions in virion and vRNA. (A) IDR outputs and contact maps of segment interactions in virion. IDR outputs from duplicated LIGR-seq of virion with or without RNA ligation were shown (left panels). Plots with < 0.01 IDR (virion RNA ligase (+)) or with < 0.05 IDR (virion RNA ligase (-)) were extracted, and contact maps of intra and intersegment interactions in virion with or without RNA ligation were plotted (middle panels). To eliminate the background interactions, we set to 0 for bins where interactions were detected with RNA ligation (-). This operation is defined as subtraction. The contract map after subtraction was shown (right panel). (B) IDR outputs and contact maps of segment interactions in vRNA. IDR outputs from duplicated LIGR-seq of vRNA with or without RNA ligation were shown (left panels). Plots with < 0.05 IDR were extracted, and contact maps of intra and intersegment interactions of vRNA with or without RNA ligation were plotted (middle panels). The contact map after subtraction was shown (right panel).

**Figure S3.**
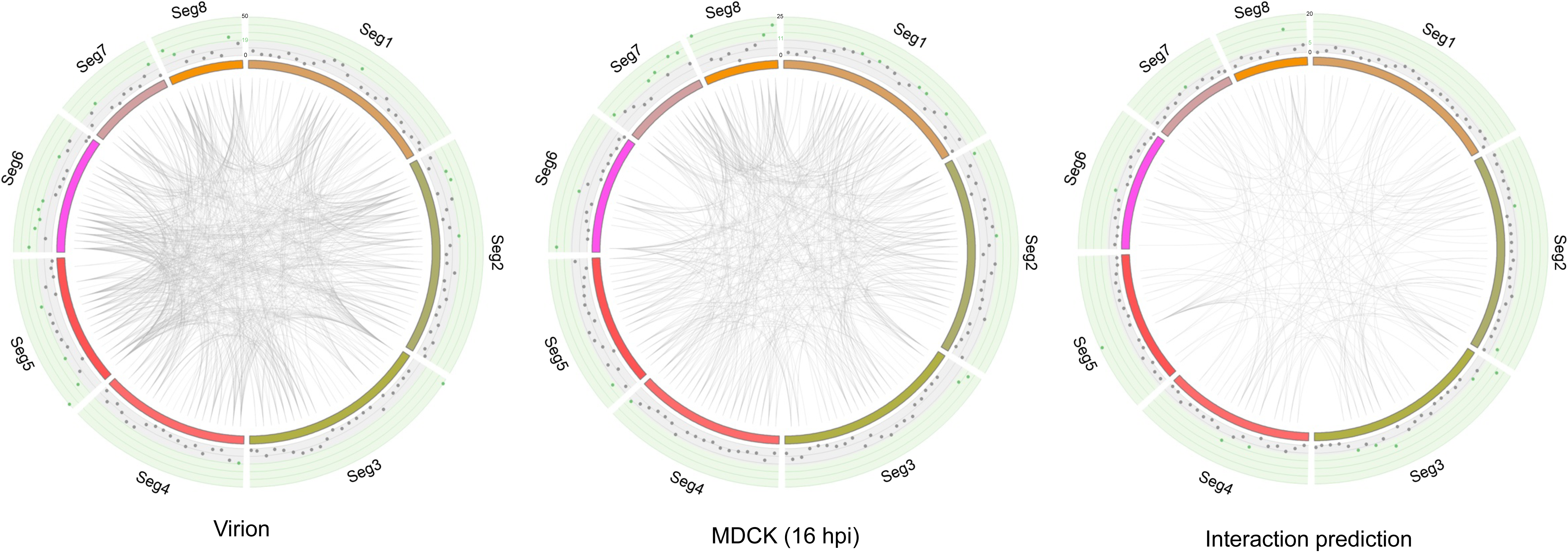
Distribution of intersegment interactions in the virion, infected cells and RNA interaction prediction. The number of intersegment interactions in each 100nt bin is plotted outside of the intersegment interaction circos plot of the virion, infected cells (16 hpi), and RNA interaction prediction. Plots of intersegment interactions over average + 1SD are shown in green.

**Figure S4.**
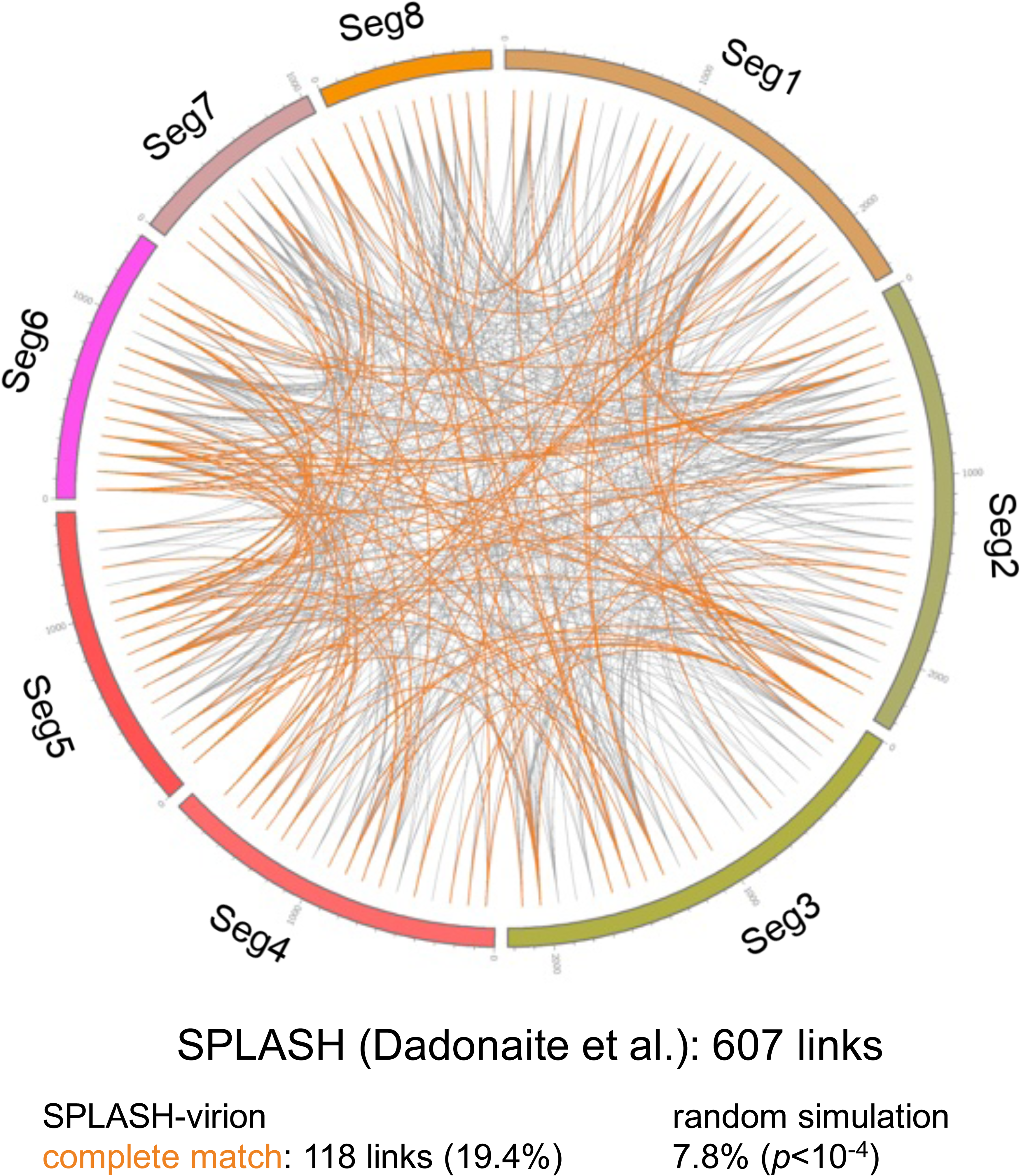
Overlapping intersegment interactions identified by customized LIGR-seq and SPLASH. Circos plot of the intersegment interactions from published SPLASH result. Intersegment interactions identified in both customized LIGR-seq and SPLASH are indicated by orange lines. “Random simulation” means the number of overlapping intersegment interactions between two randomly generated intersegment interaction sets.

**Figure S5.**
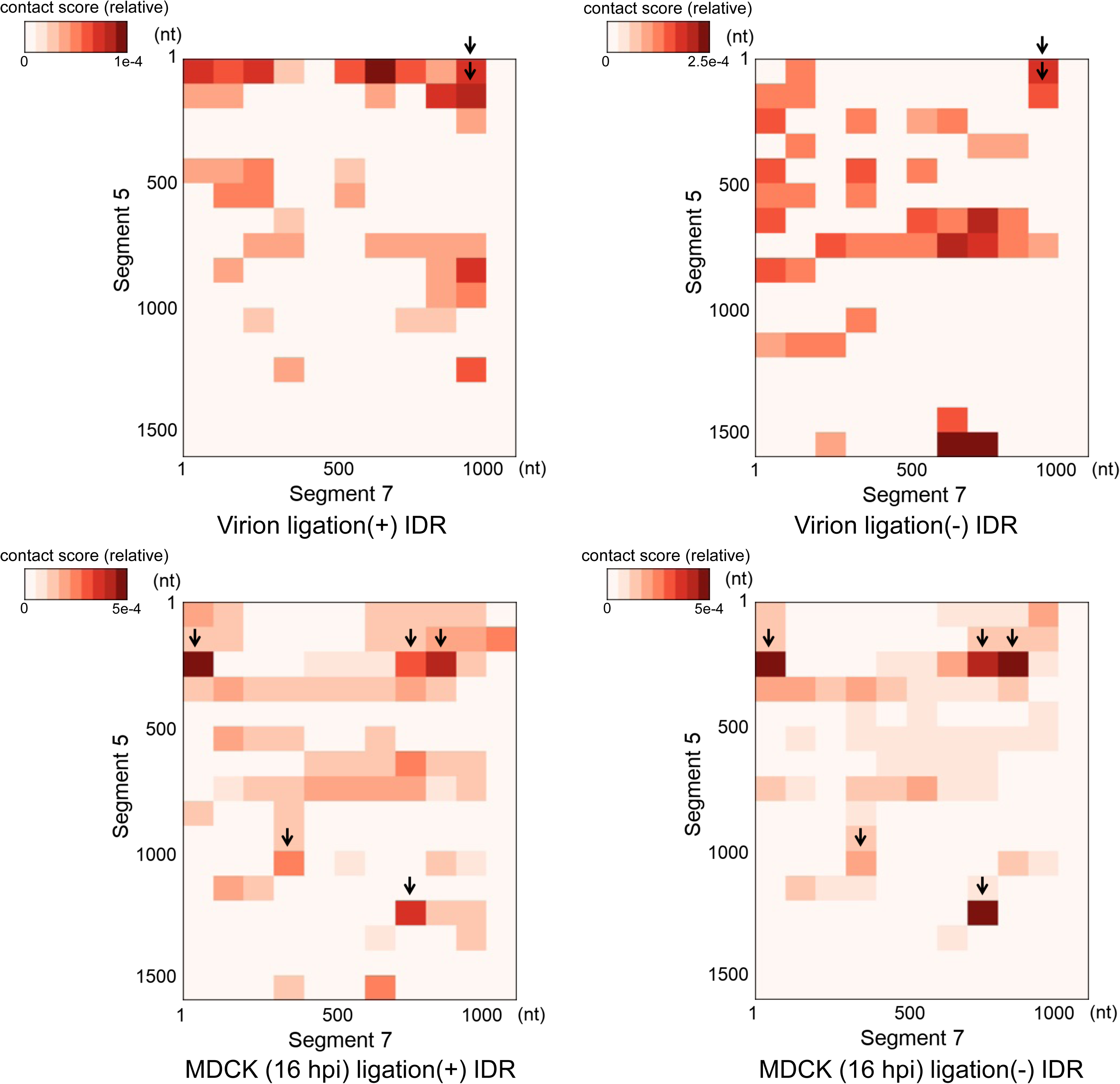
Intersegment interactions between segments 5 and 7 identified both with and without RNA ligation samples. Heatmaps of intersegment interactions between segment 5 and 7 in the virion and infected cells (16 hpi) with and without RNA ligation are shown. Arrows indicate bins that show strong interaction signals both with and without RNA ligation.

**Figure S6.**
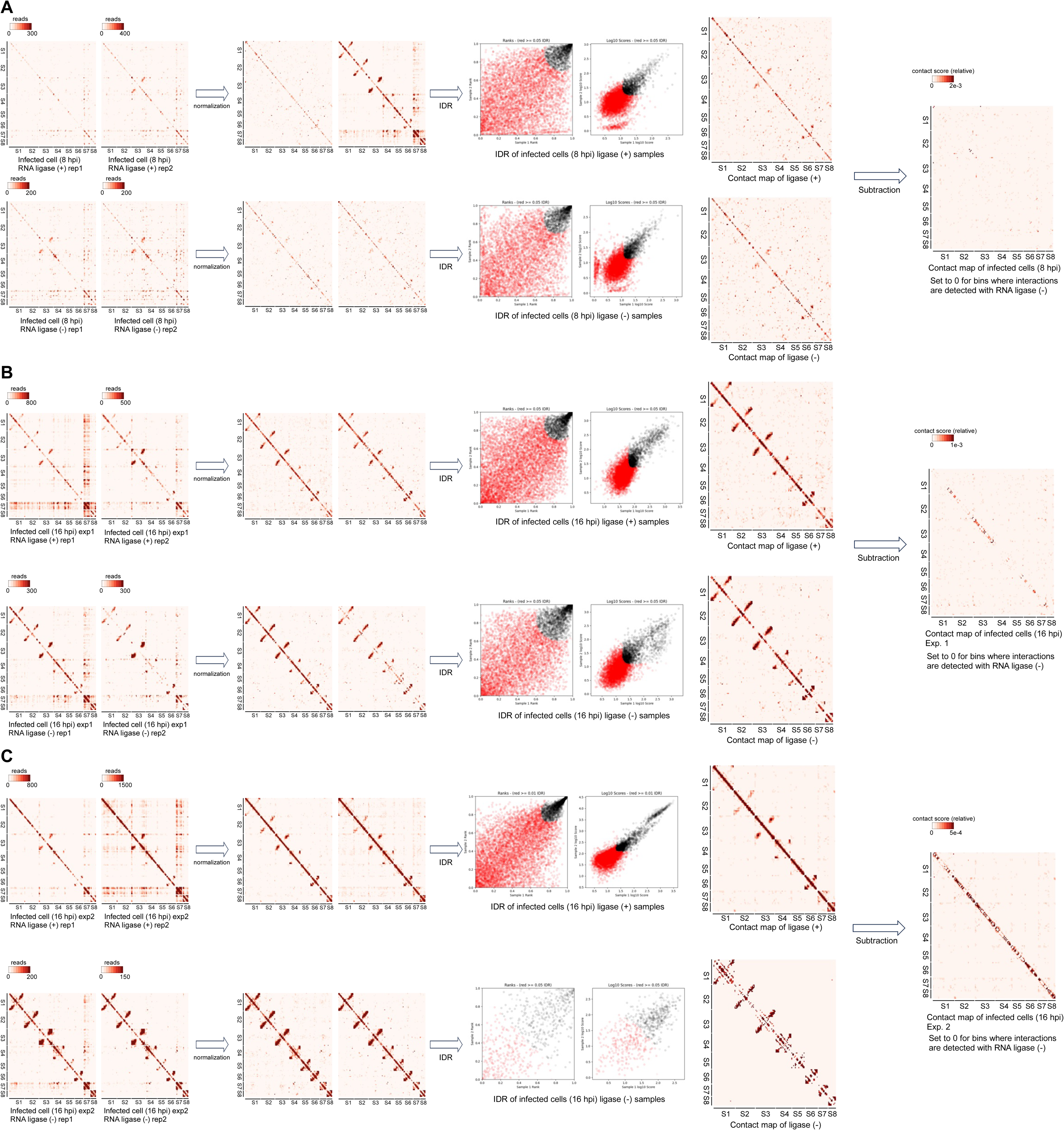
IDR outputs and contact maps of segment interactions in infected cells. (A) IDR outputs and contact maps of segment interactions in infected cells at 8 hpi. IDR outputs and contact maps before and after subtraction were shown. (B and C) IDR outputs and contact maps of segment interactions in infected cells at 16 hpi. The results of two independent experiments are shown in (B) and (C). IDR outputs and contact maps before and after subtraction were shown.

**Figure S7.**
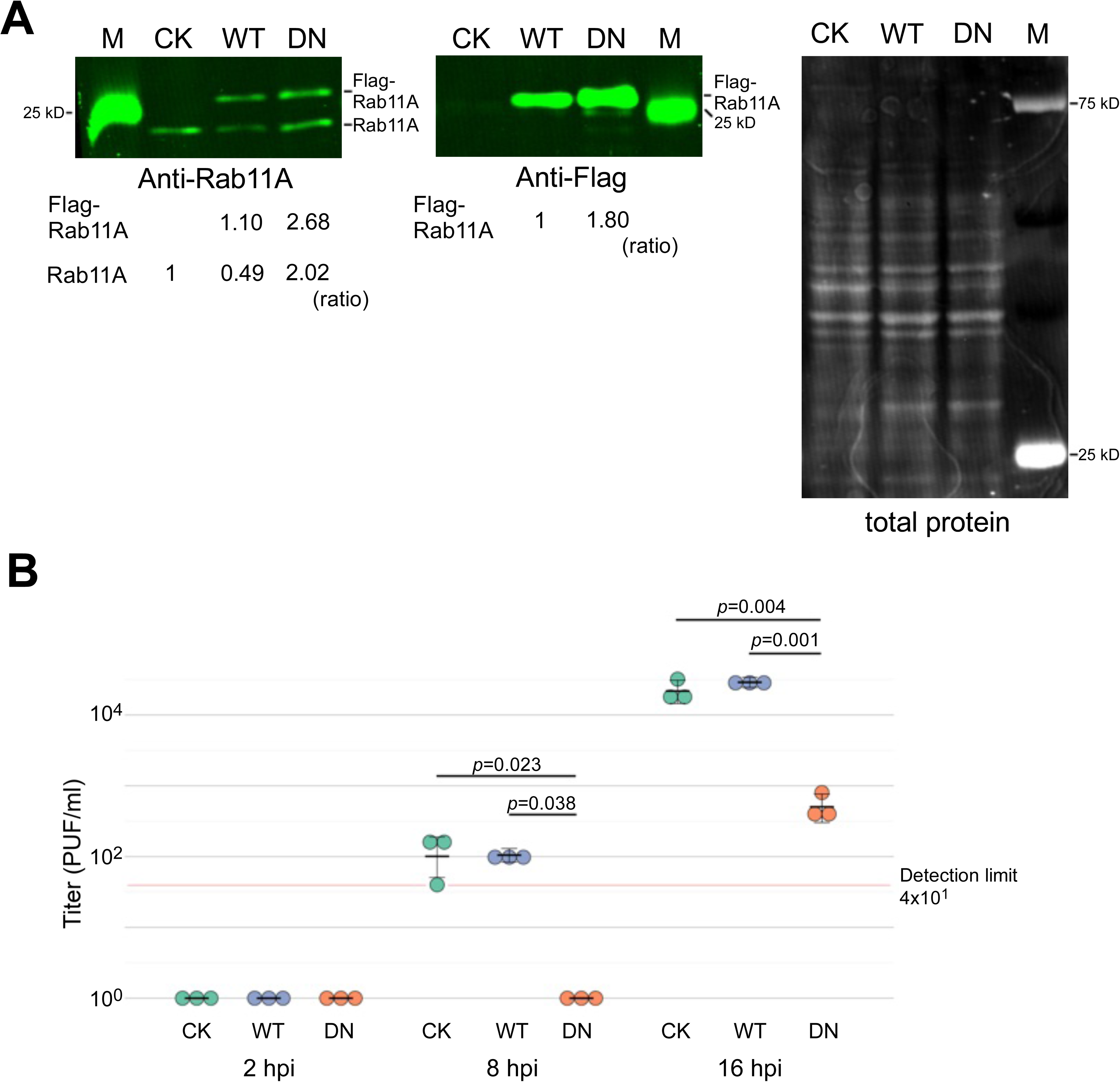
Expression of Flag-Rab11 and virus propagation in MDCK-Flag Rab11wt and MDCK-Flag Rab11DN cells. (A) Quantification of Rab11A in MDCK, MDCK-Flag Rab11wt, and MDCK-Flag Rab11DN by fluorescence western blotting. Endogenous Rab11A and Flag-Rab11A were stained by anti-Rab11A and anti-Flag antibodies. Total protein was stained by Revert 520 Total Protein Stain Solution, and the amount of total protein was used for normalization. CK: MDCK, WT: MDCK-Flag Rab11wt, DN: MDCK-Flag Rab11DN. (B) Virus titer in the supernatant from infected MDCK, MDCK-Flag Rab11wt, and MDCK-Flag Rab11DN cells. Virus titer at 2 hpi, 8 hpi, and 16 hpi was determined by plaque assay. The crossbars indicate average values with standard deviations from three independent experiments. The circles indicate the titer of each experiment. The red line indicates the lower limit of detection of virus titer. P-values were calculated by Dunnett’s multiple comparison test. CK: MDCK, WT: MDCK-Flag Rab11wt, DN: MDCK-Flag Rab11DN.

**Figure S8.**
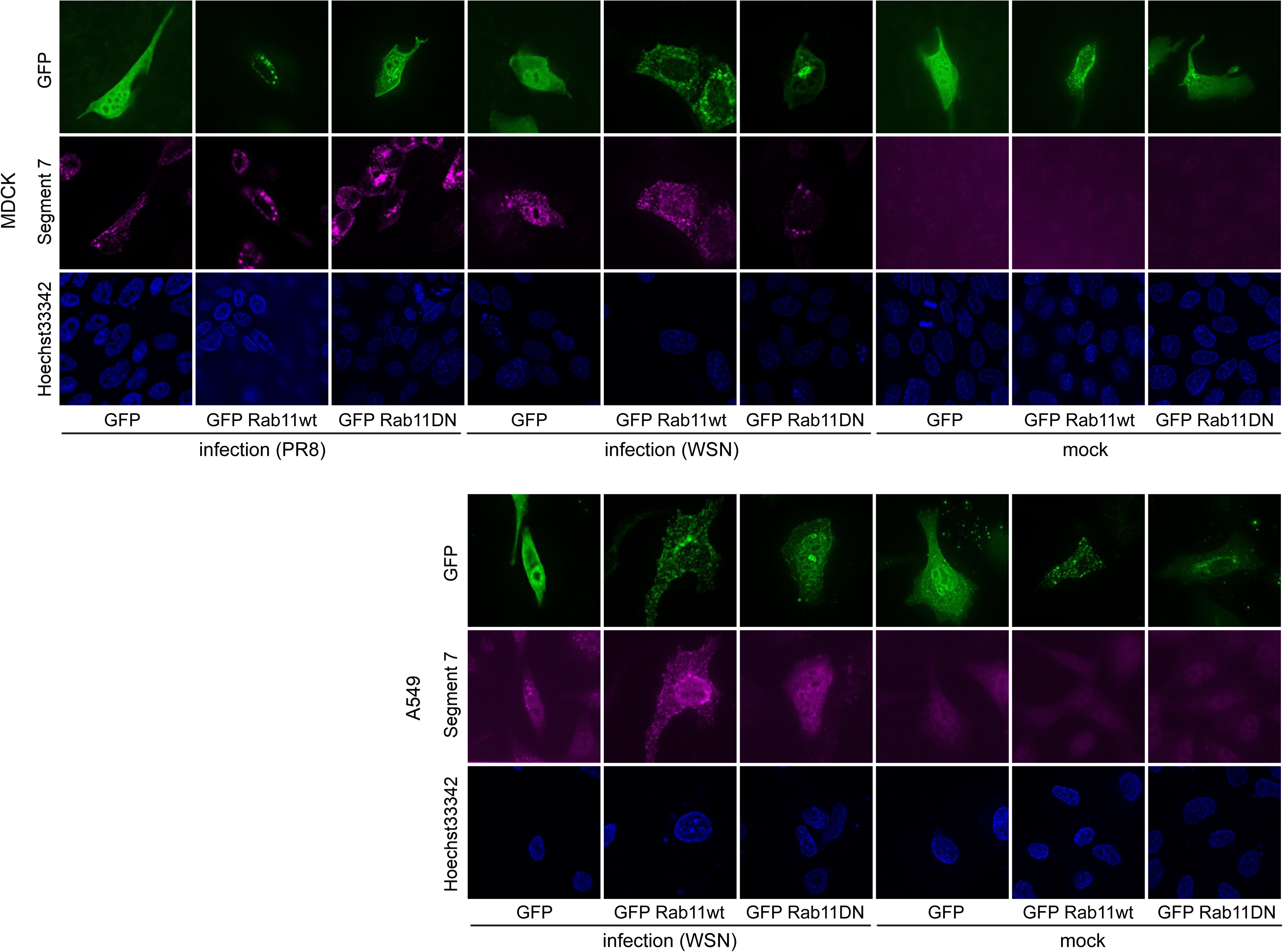
Localization of vRNA in MDCK and A549 cells transiently expressing GFP-Rab11DN. MDCK and A549 cells were infected with PR8 or WSN at an MOI of 1 (MDCK) or 5 (A549). vRNA (Segment 7) was visualized by in situ hybridization at 16 hpi. Bar: 5 µm.

**Figure S9.**
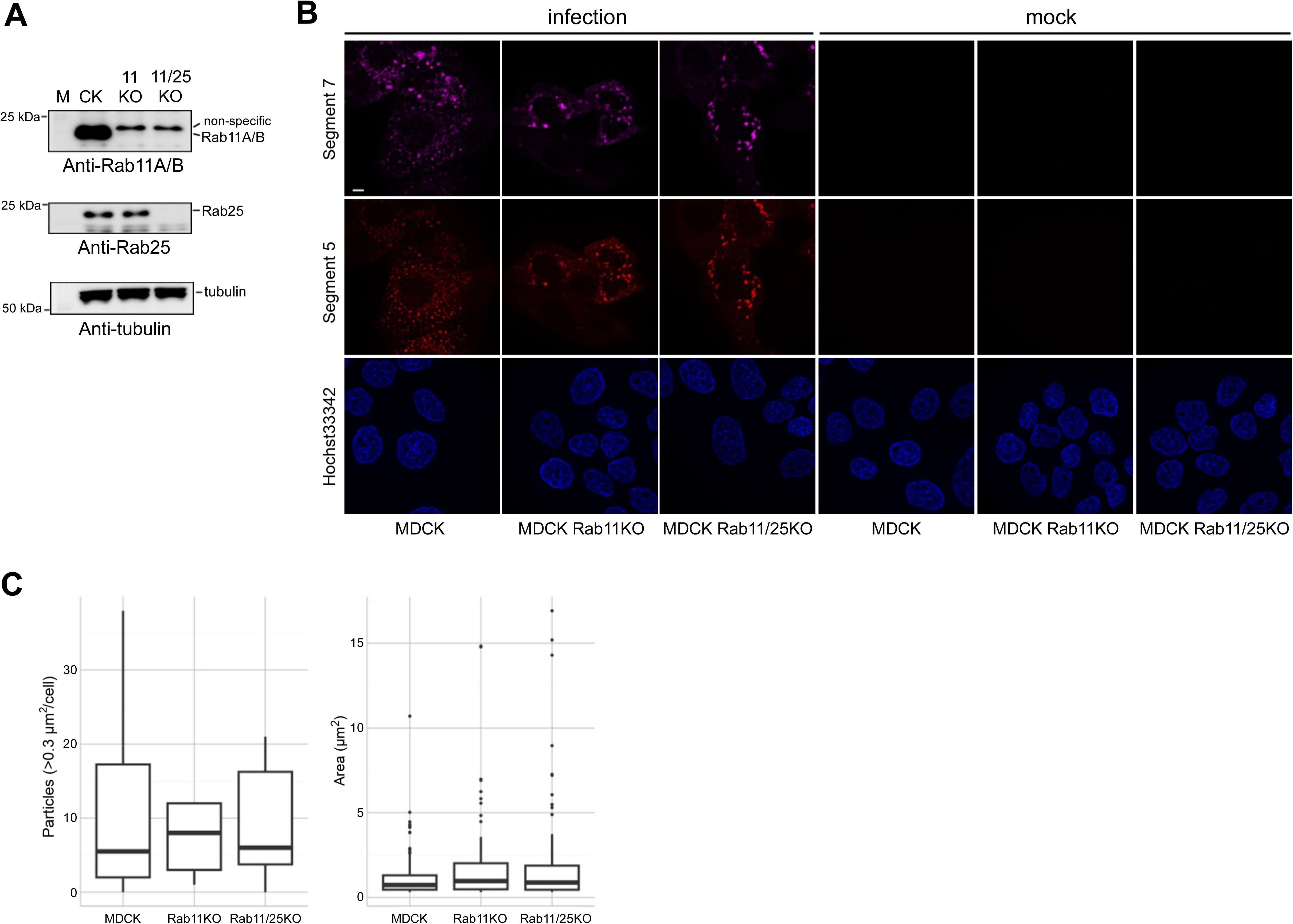
Formation of viral inclusions in Rab11KO and Rab11/25 KO cells. (A) Confirmation of knock-down of Rab11 and Rab25 in MDCK Rab11KO and MDCK Rab11/25 KO cells. Knock-down of Rab11 and Rab25 was confirmed by fluorescence western blotting. M: marker, CK: MDCK cells, 11KO: MDCK Rab11KO cells, and 11/25KO: MDCK Rab11/25KO cells. (B) Formation of viral inclusions in Rab11KO and Rab11/25KO cells. Bar: 5 µm. (C) The particle number and size of viral inclusions in infected MDCK, MDCK Rab11KO, and MDCK Rab11/25KO cells. Number of measured cells is 16 (MDCK), 9 (MDCK Rab11KO), or 12 (MDCK Rab11/25KO). Number of measured particles is 178 (MDCK), 67 (MDCK Rab11KO), or 112 (MDCK Rab11/25KO).

**Figure S10.**
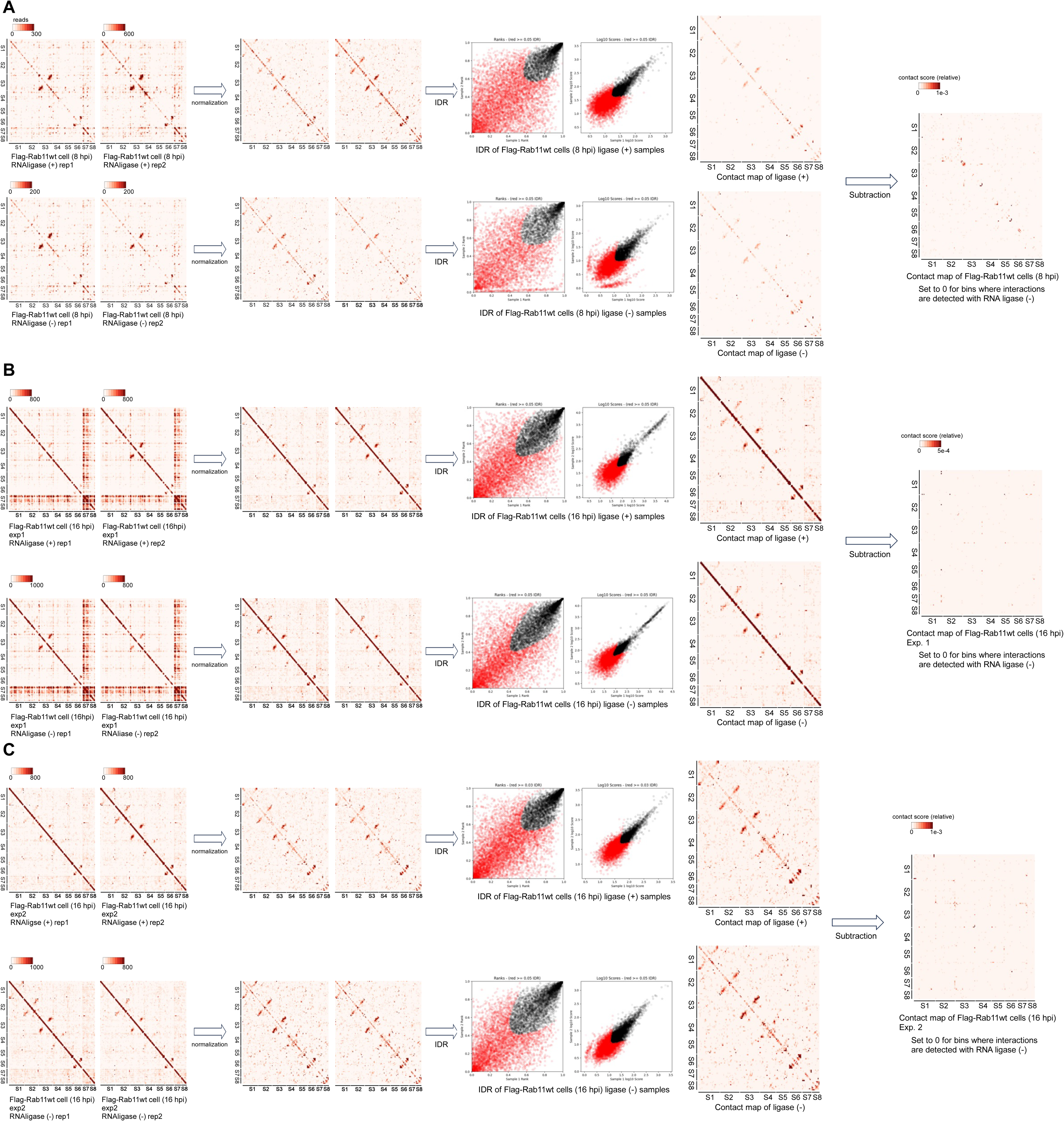
IDR outputs and contact maps of segment interactions in infected cells expressing Flag-Rab11wt. (A-C) IDR outputs and contact maps of segment interactions in cells expressing Flag-Rab11wt at 8 hpi (A) and 16 hpi (B and C). The results of two independent experiments are shown in (B) and (C).

**Figure S11.**
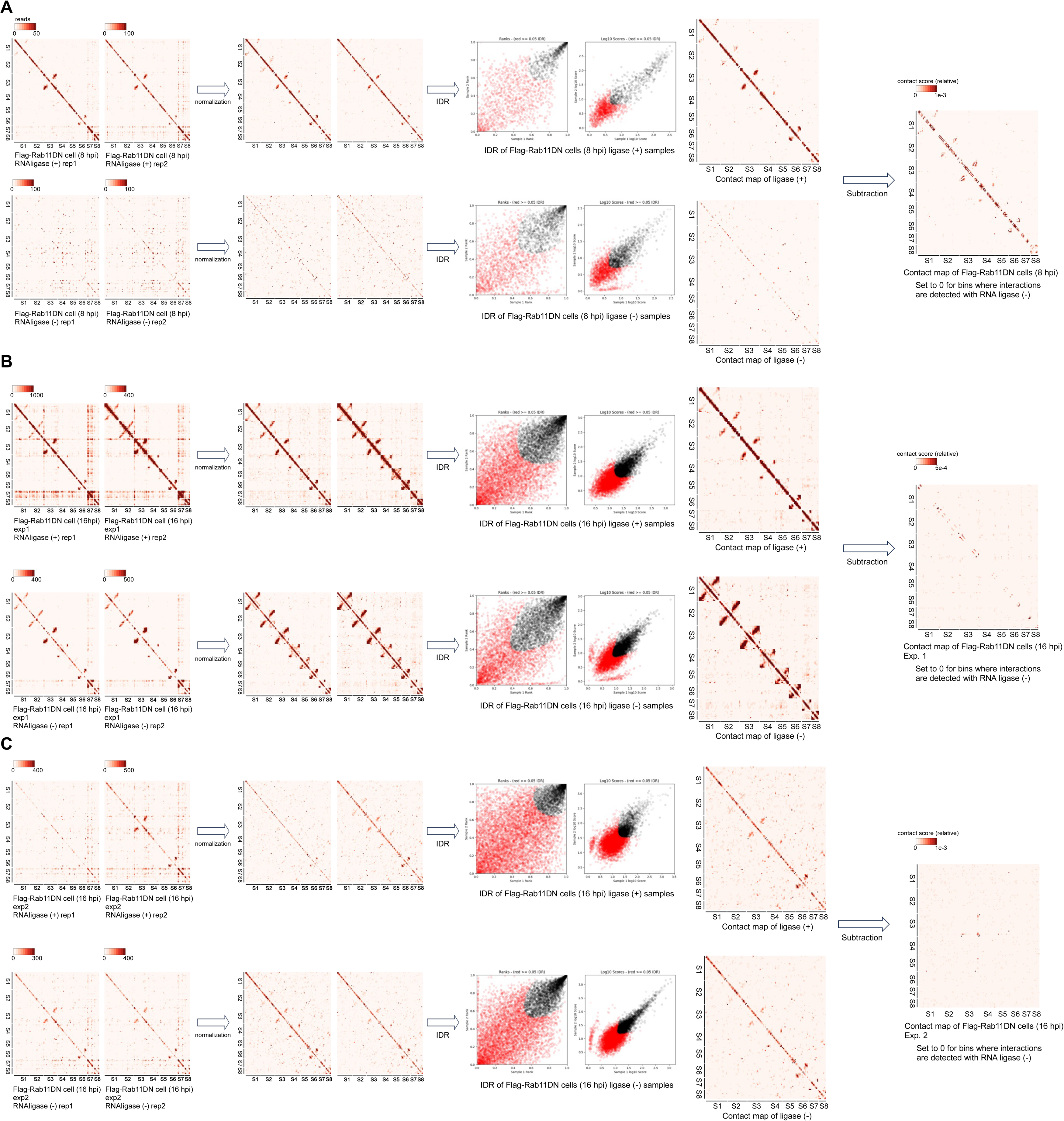
IDR outputs and contact maps of segment interactions in infected cells expressing Flag-Rab11DN. (A-C) IDR outputs and contact maps of segment interactions in cells expressing Flag-Rab11DN at 8 hpi (A) and 16 hpi (B and C). The results of two independent experiments are shown in (B) and (C).

**Figure S12.**
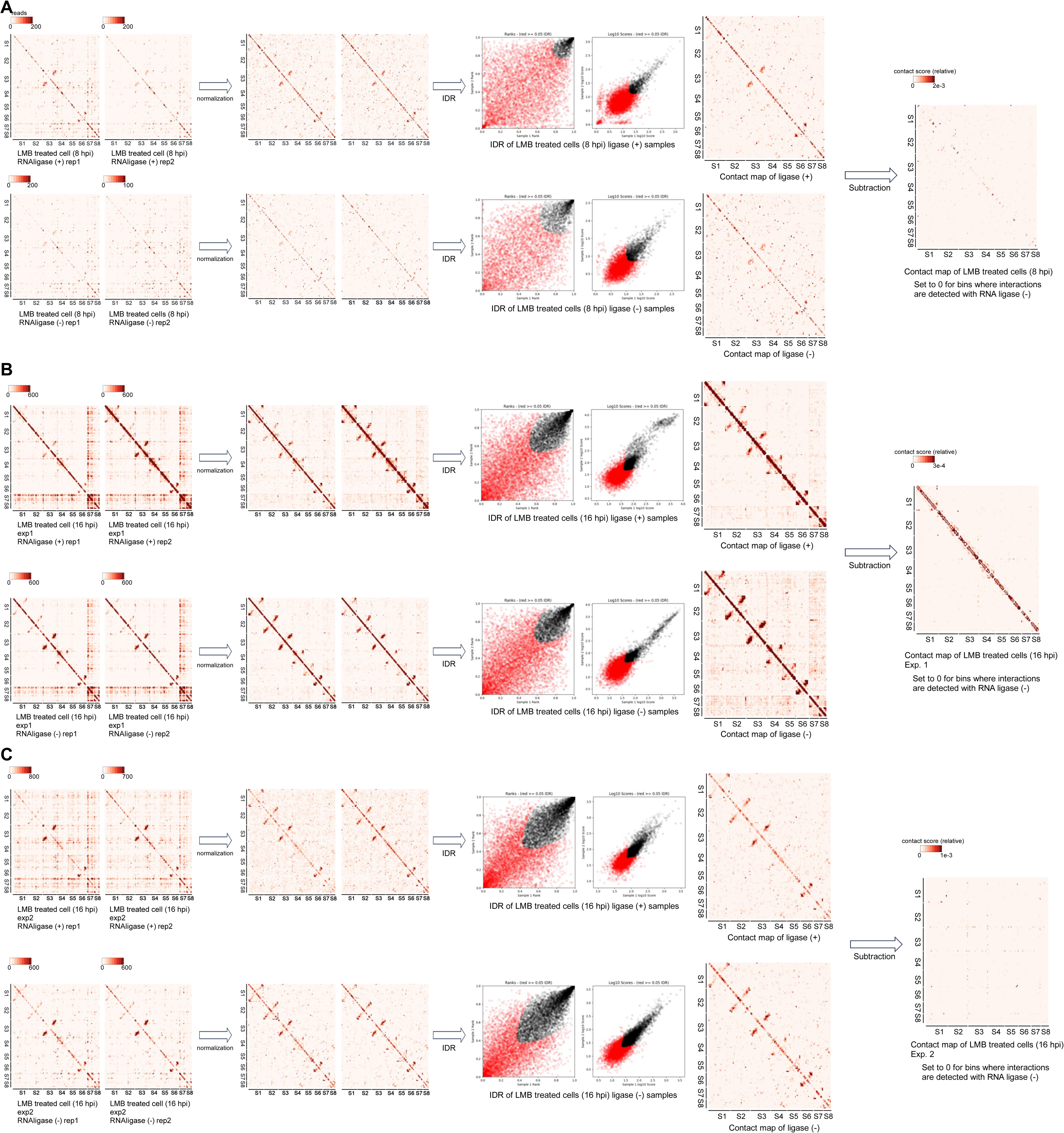
IDR outputs and contact maps of segment interactions in infected cells with leptomycin B treatment. (A) IDR outputs and contact maps of segment interactions in infected cells with leptomycin B (LMB) treatment at 8 hpi. (B and C) IDR outputs and contact maps of segment interactions in infected cells with LMB treatment at 16 hpi. The results of two independent experiments are shown in (B) and (C).

**Video S1, S2, and S3.** PA-GFP movement in MDCK (S1), MDCK-Flag Rab11wt (S2), and MDCK-Flag Rab11DN (S3).

**Video S4, S5, and S6.** FRAP experiments of PA-GFP in MDCK (S4), MDCK-Flag Rab11wt (S5), and MDCK-Flag Rab11DN (S6).

**Supplemental Table.**
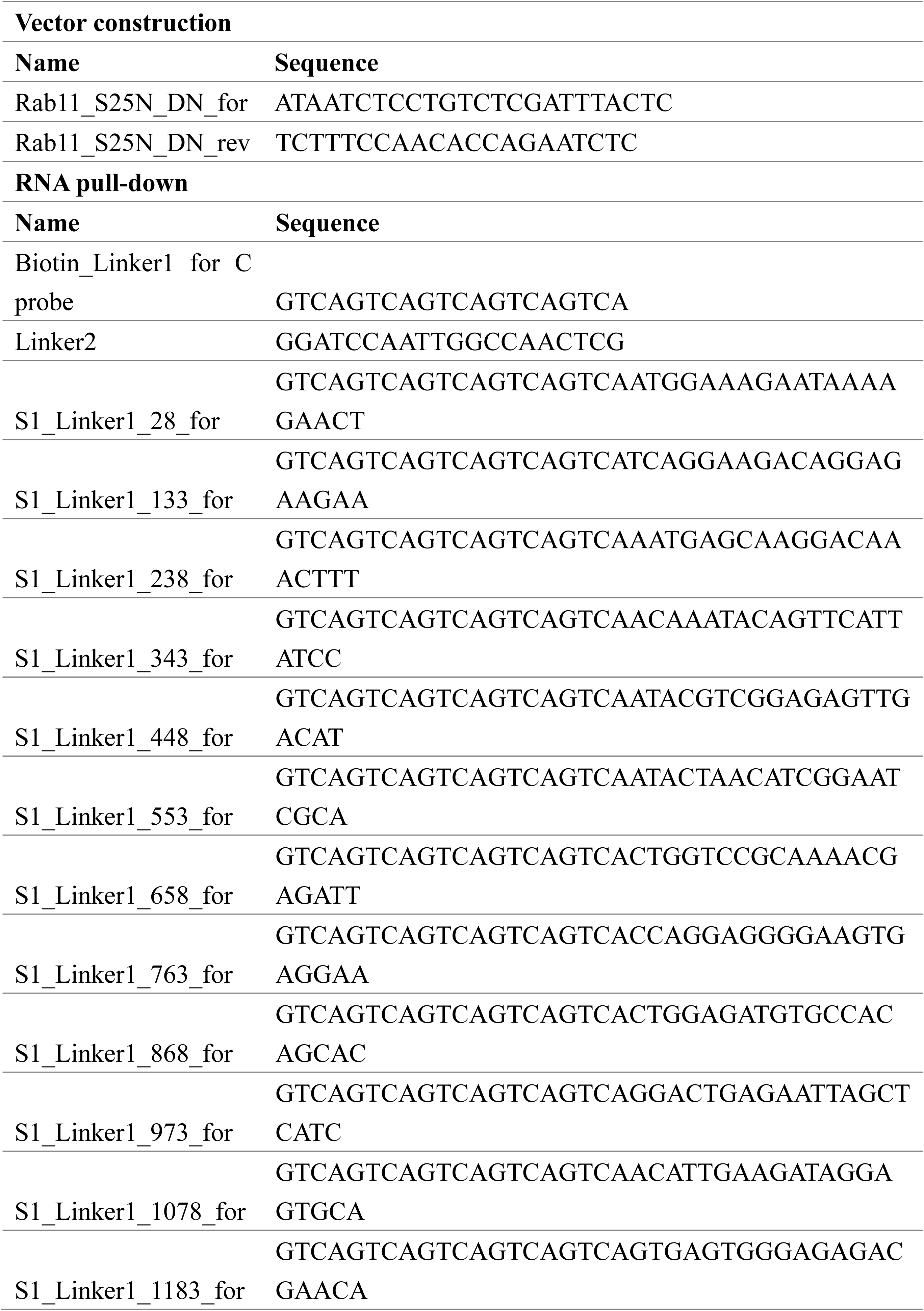
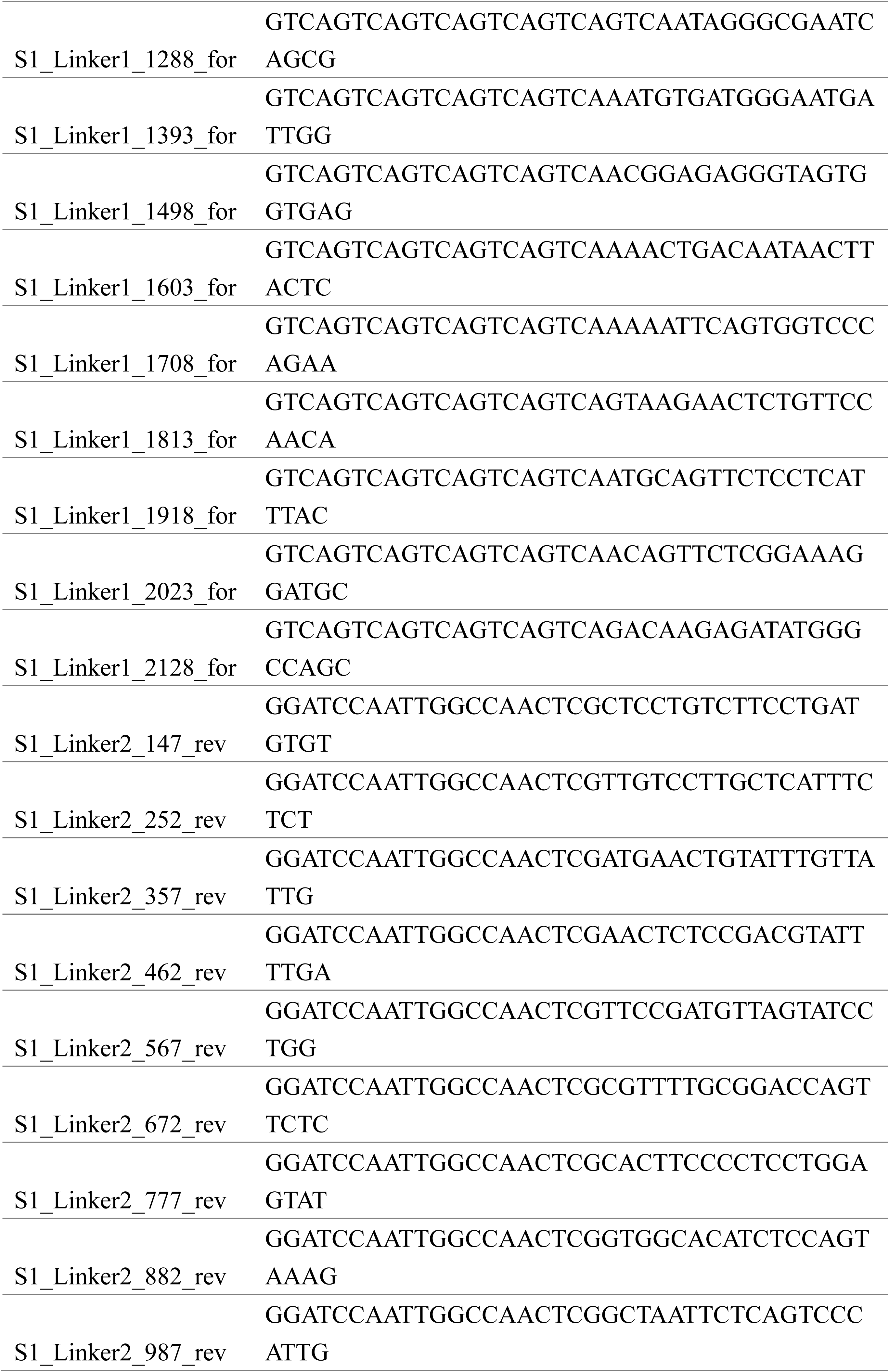
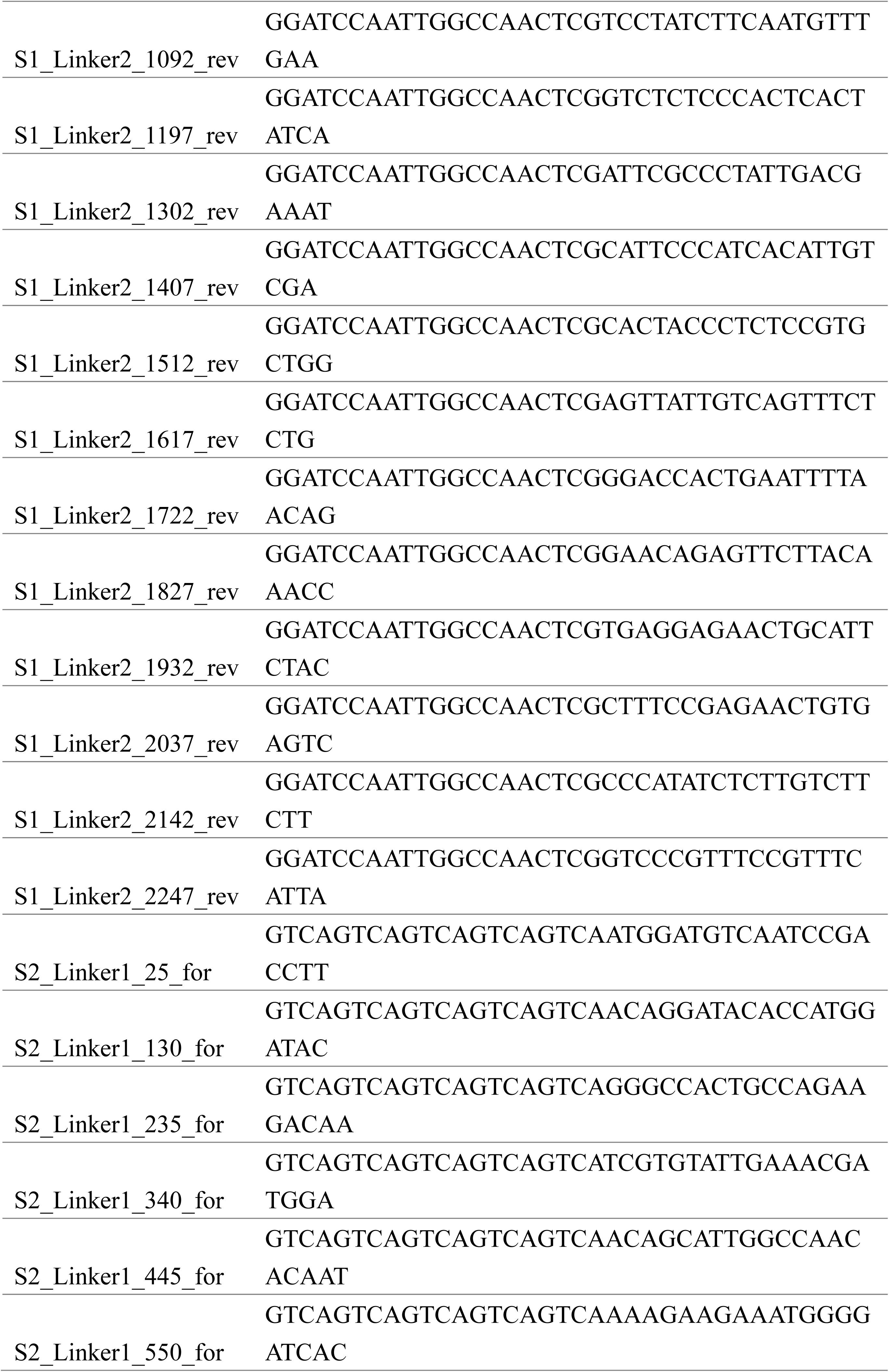
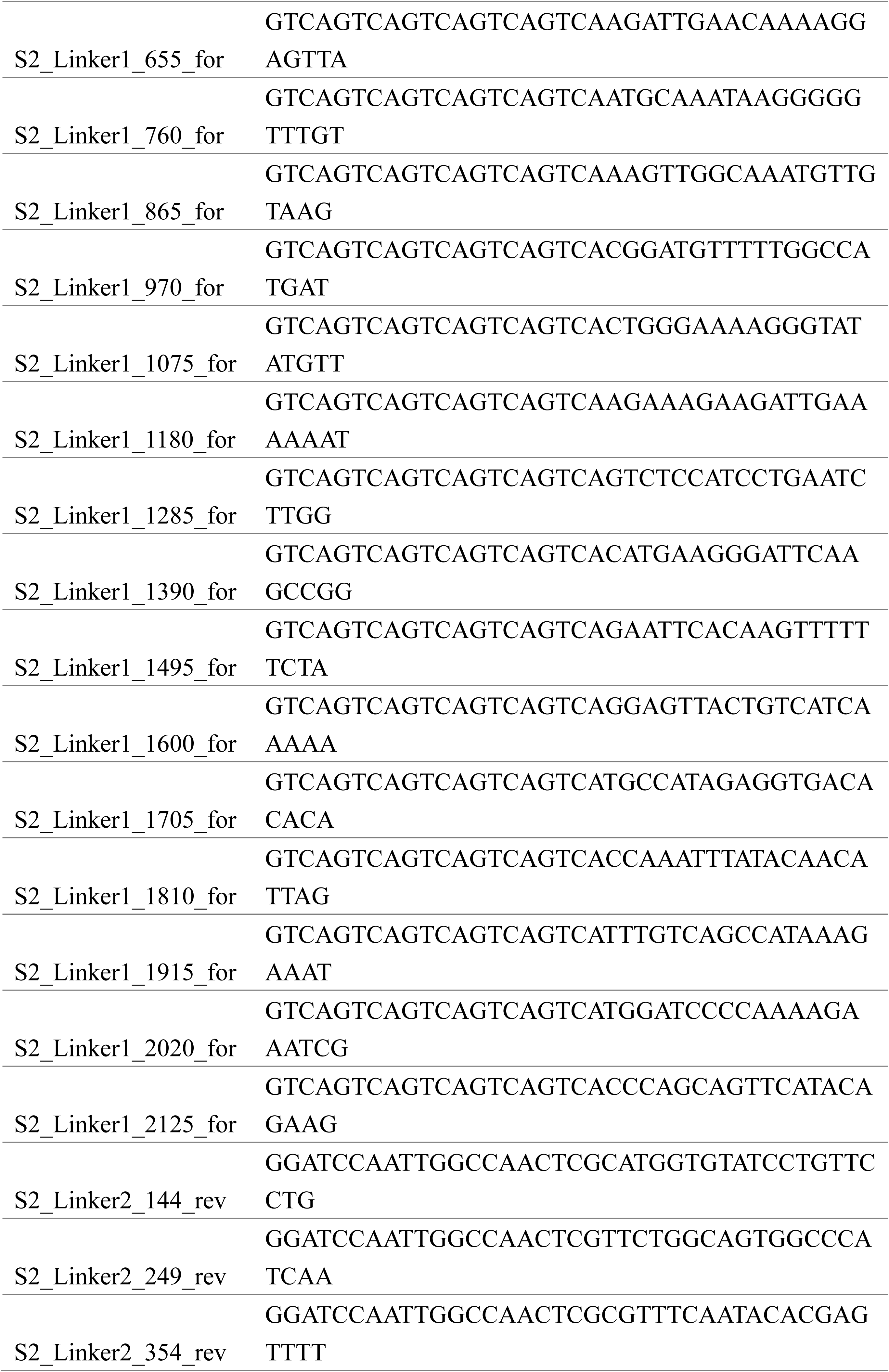
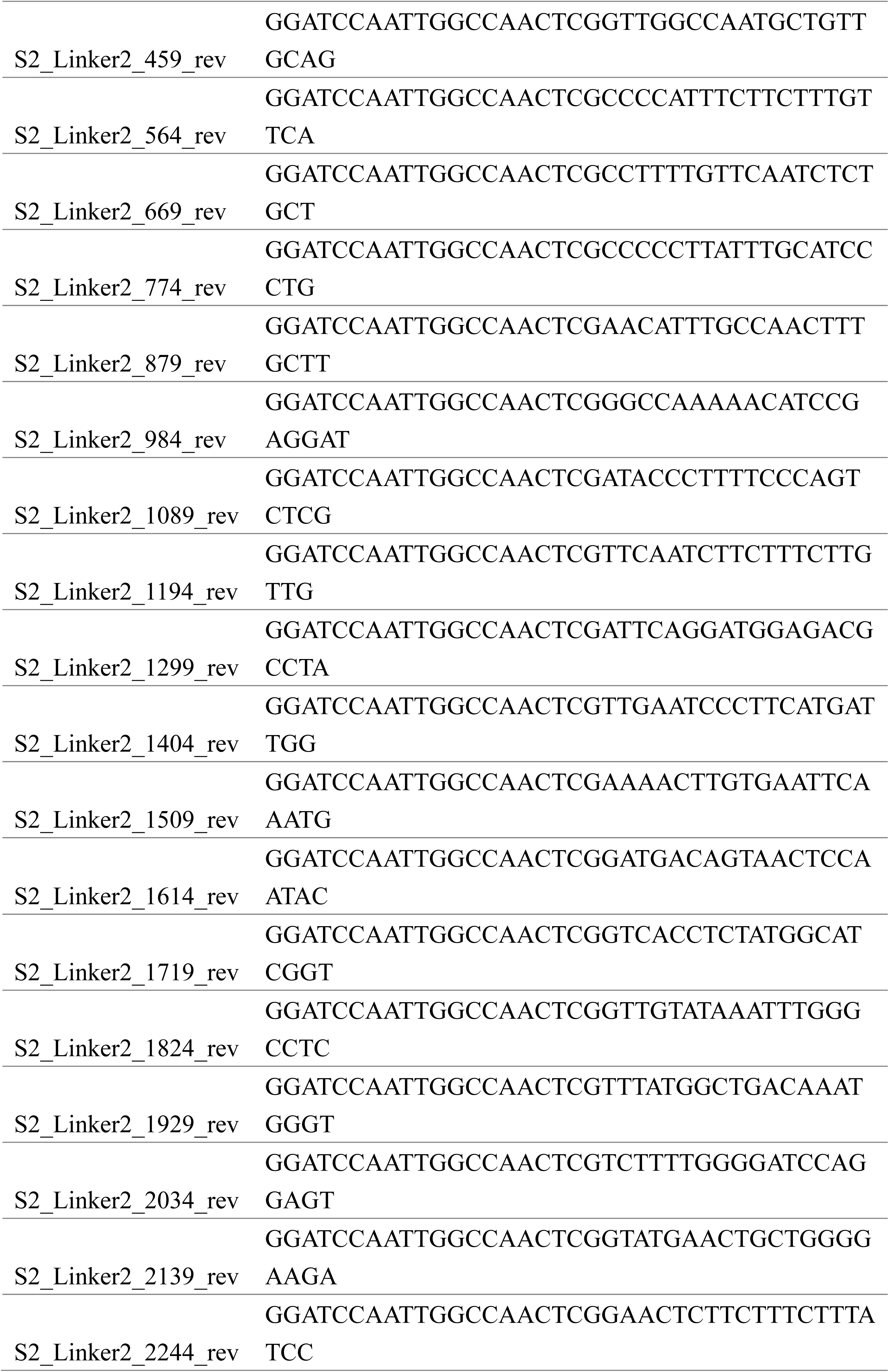
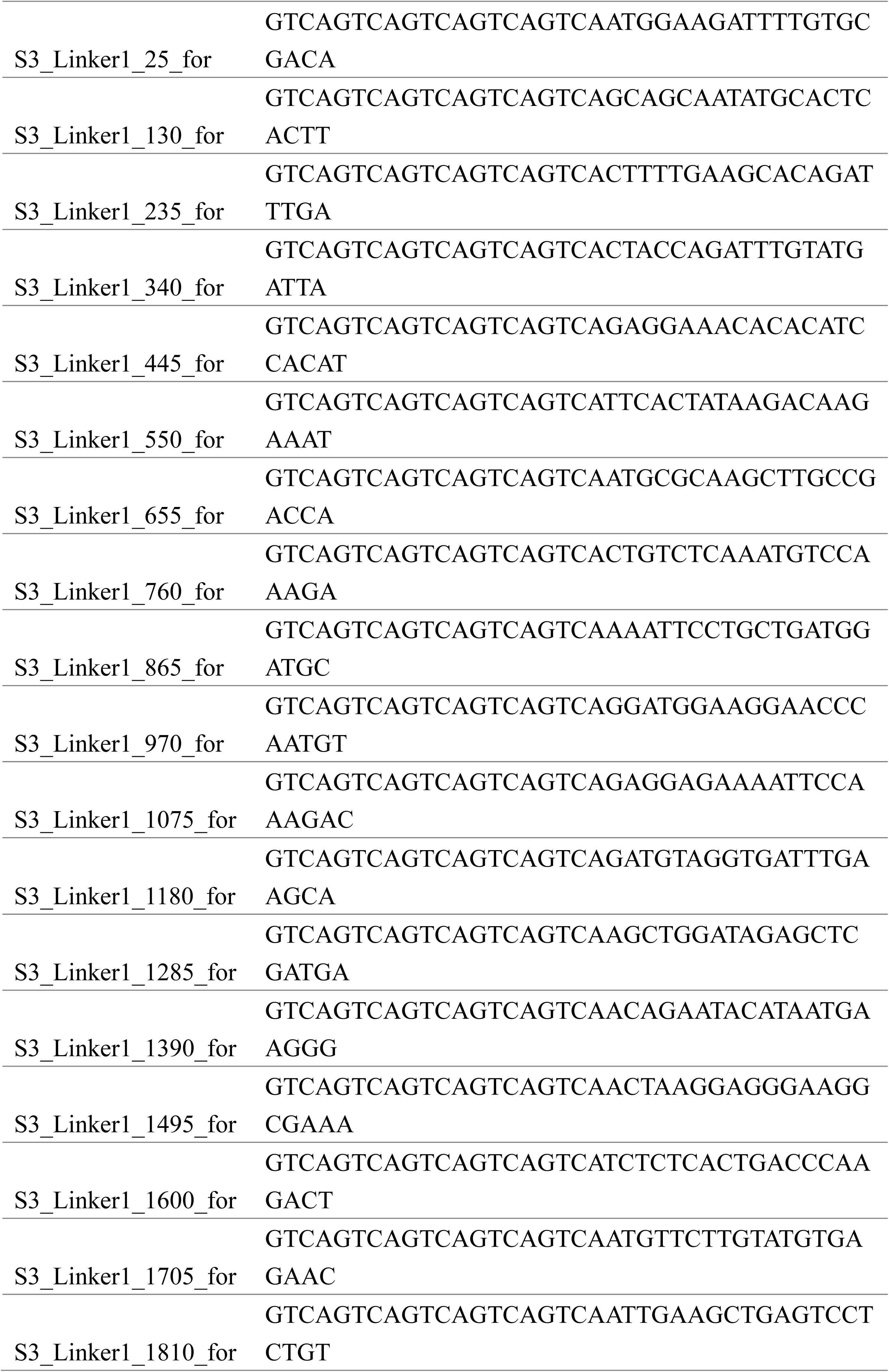
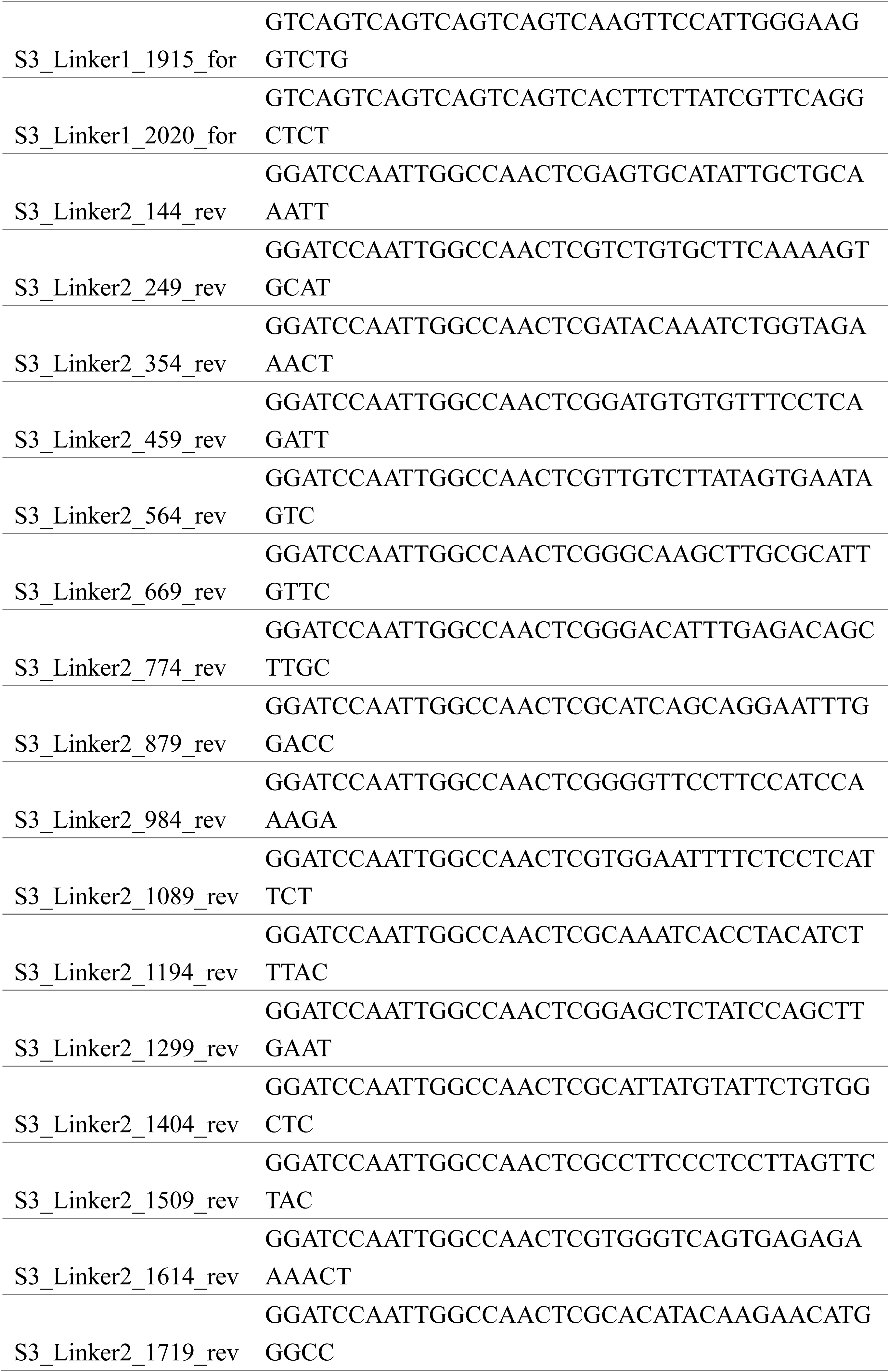
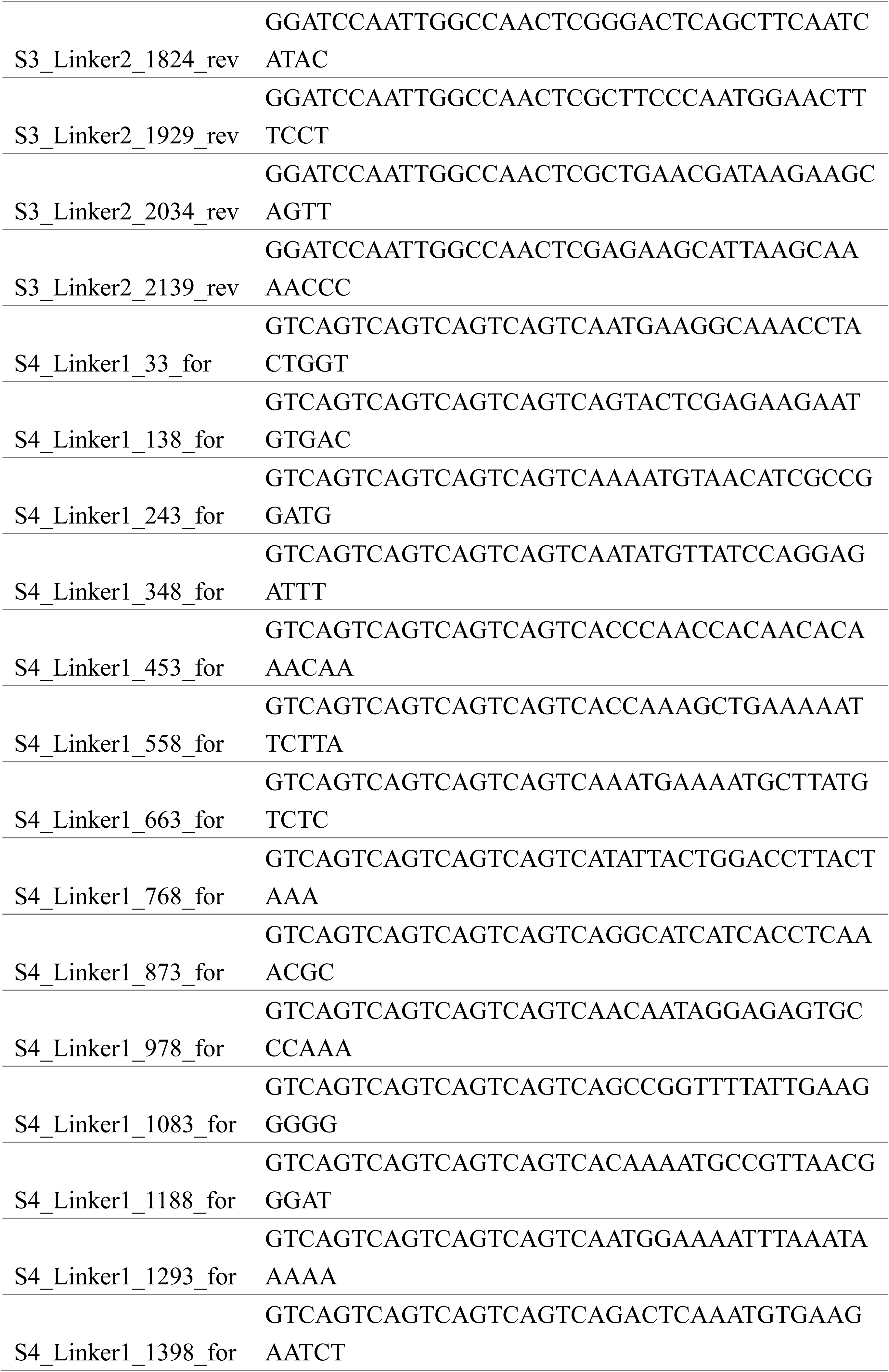
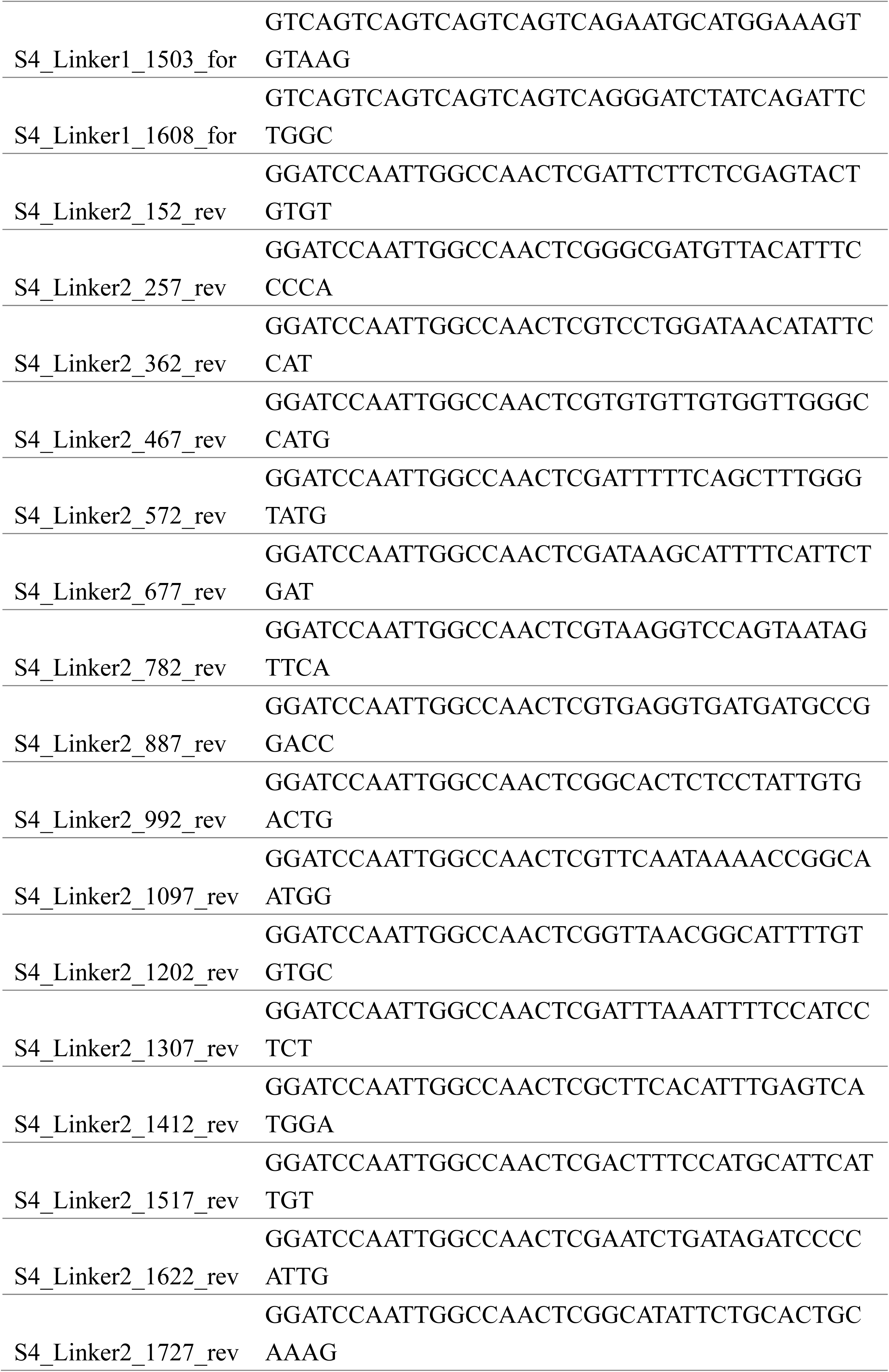
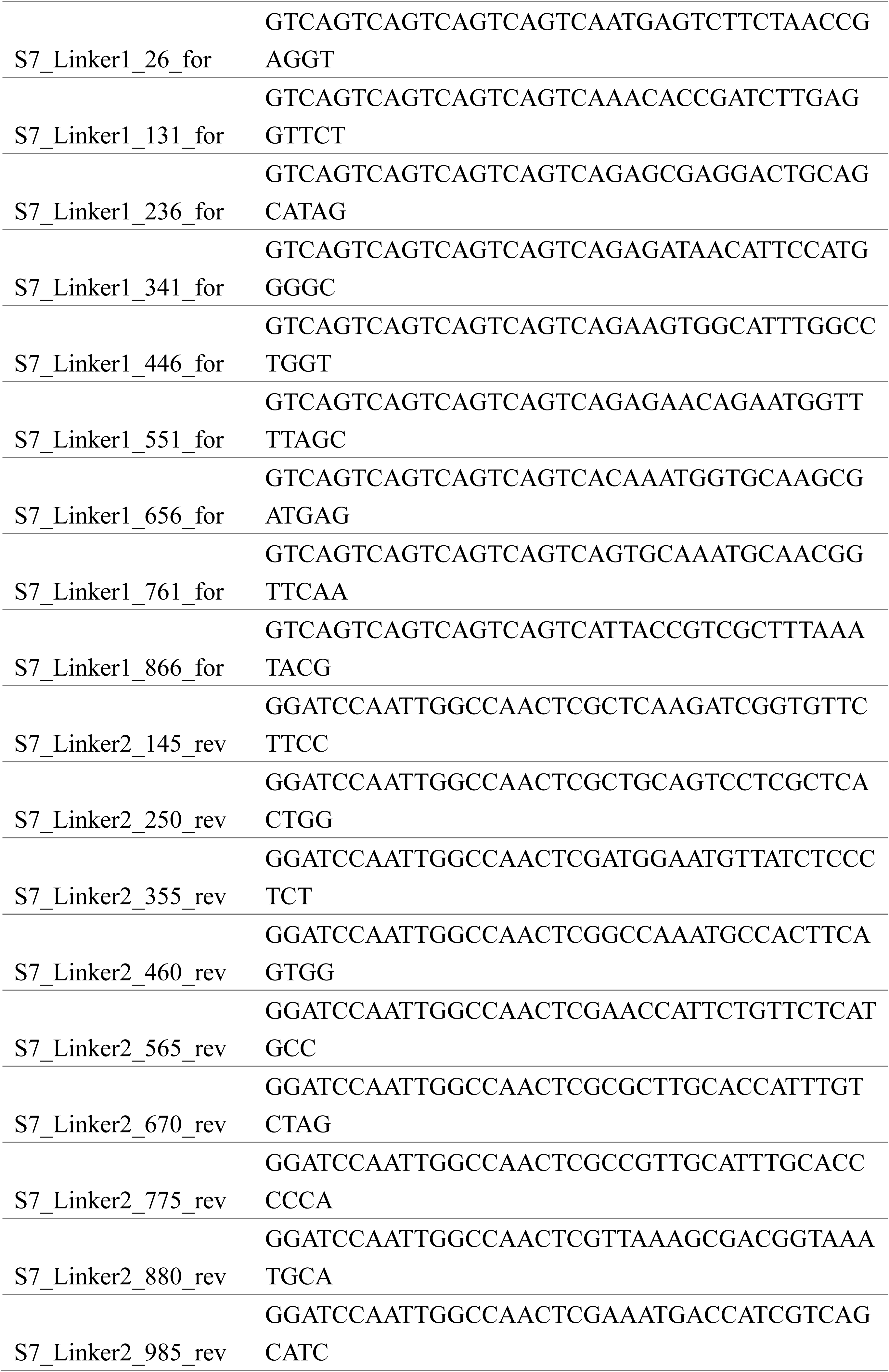
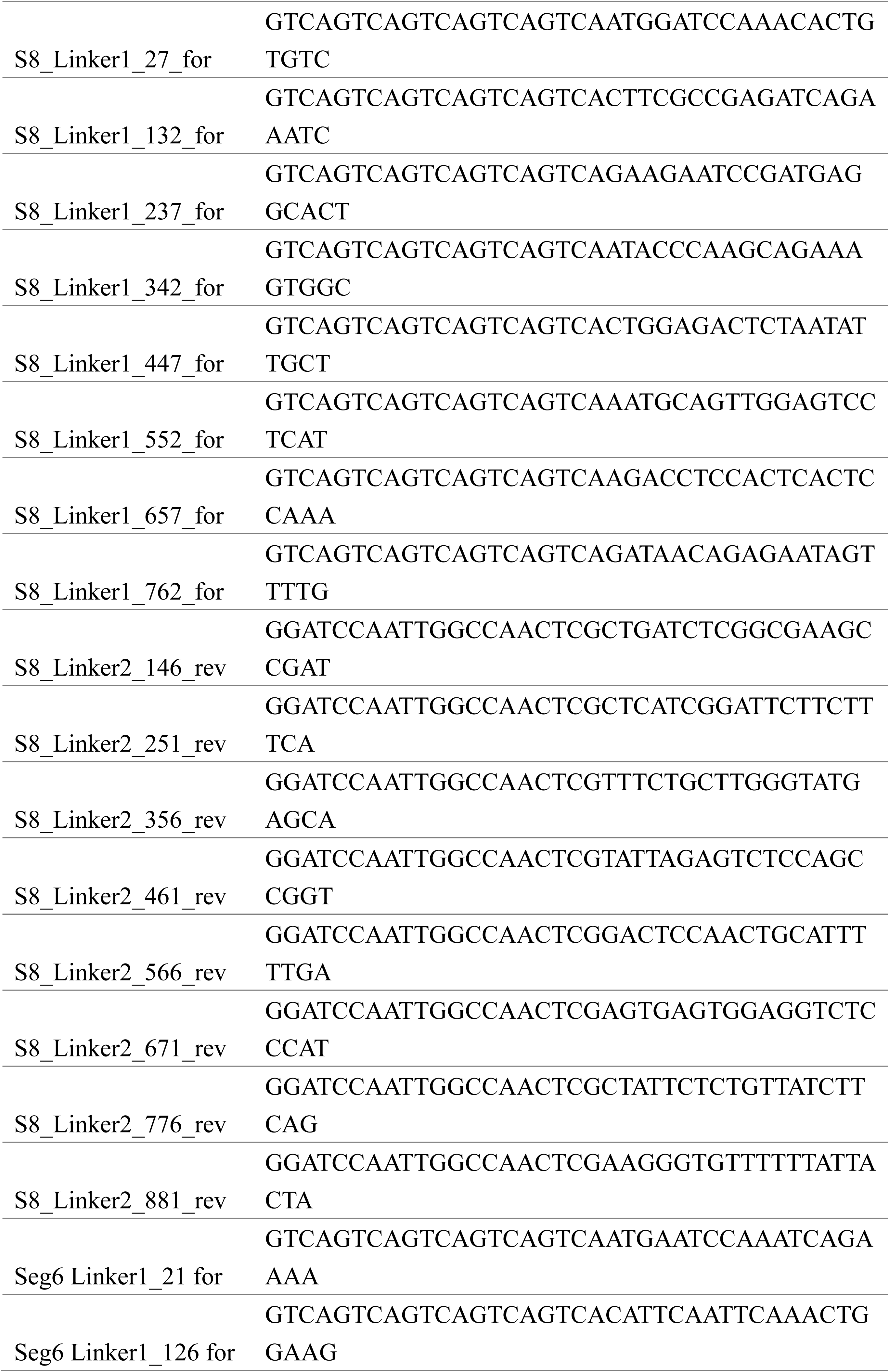
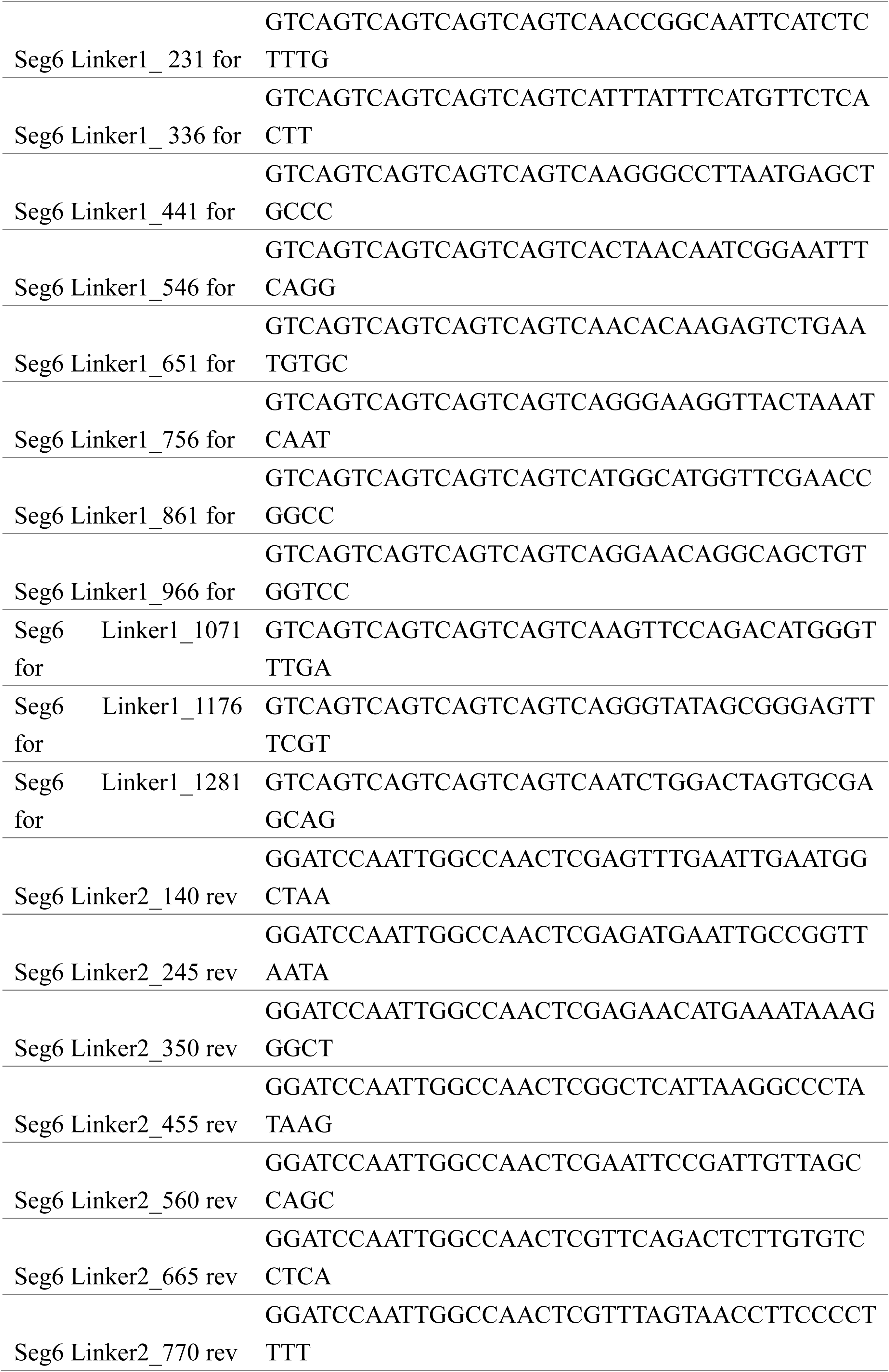
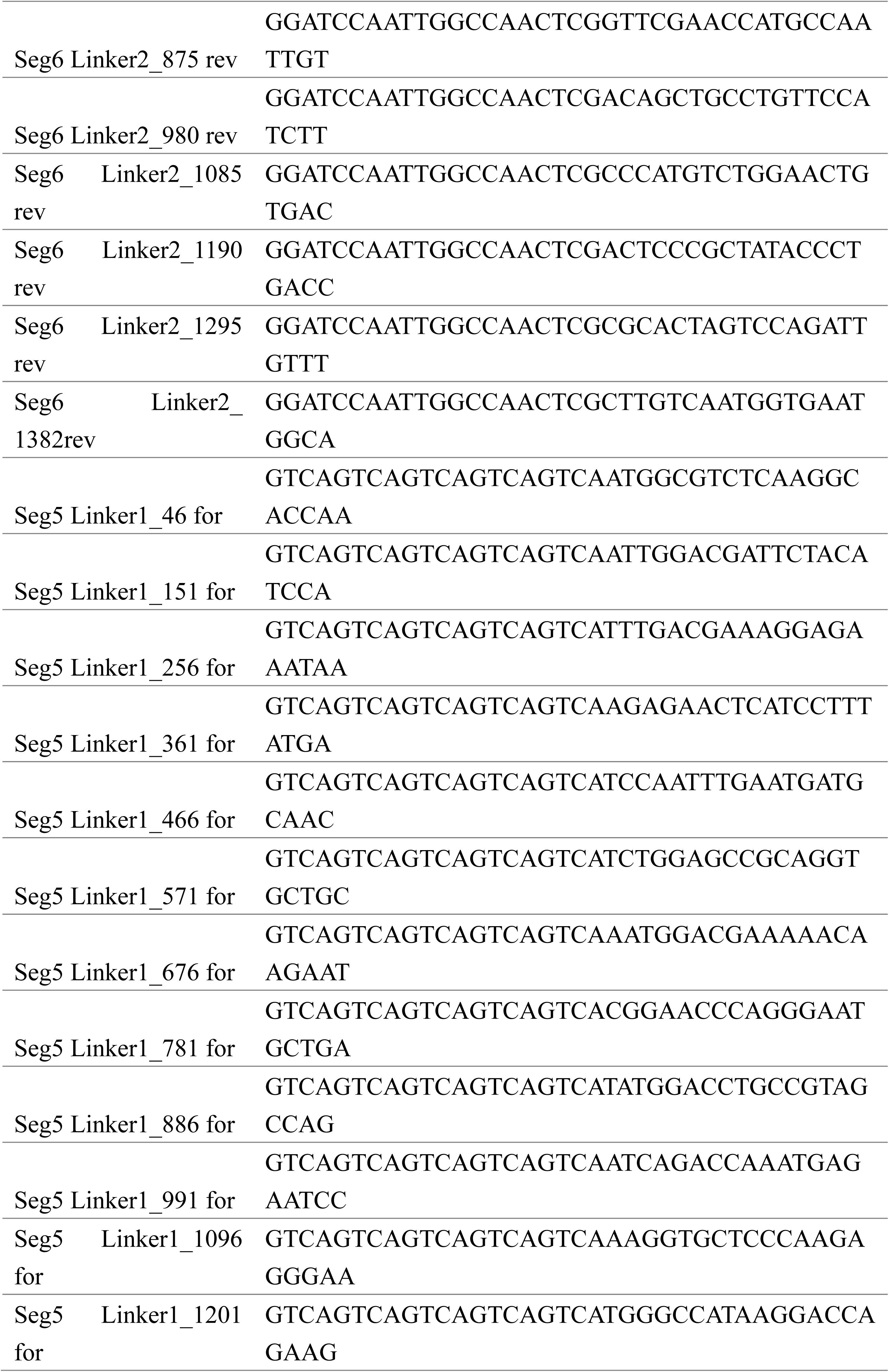
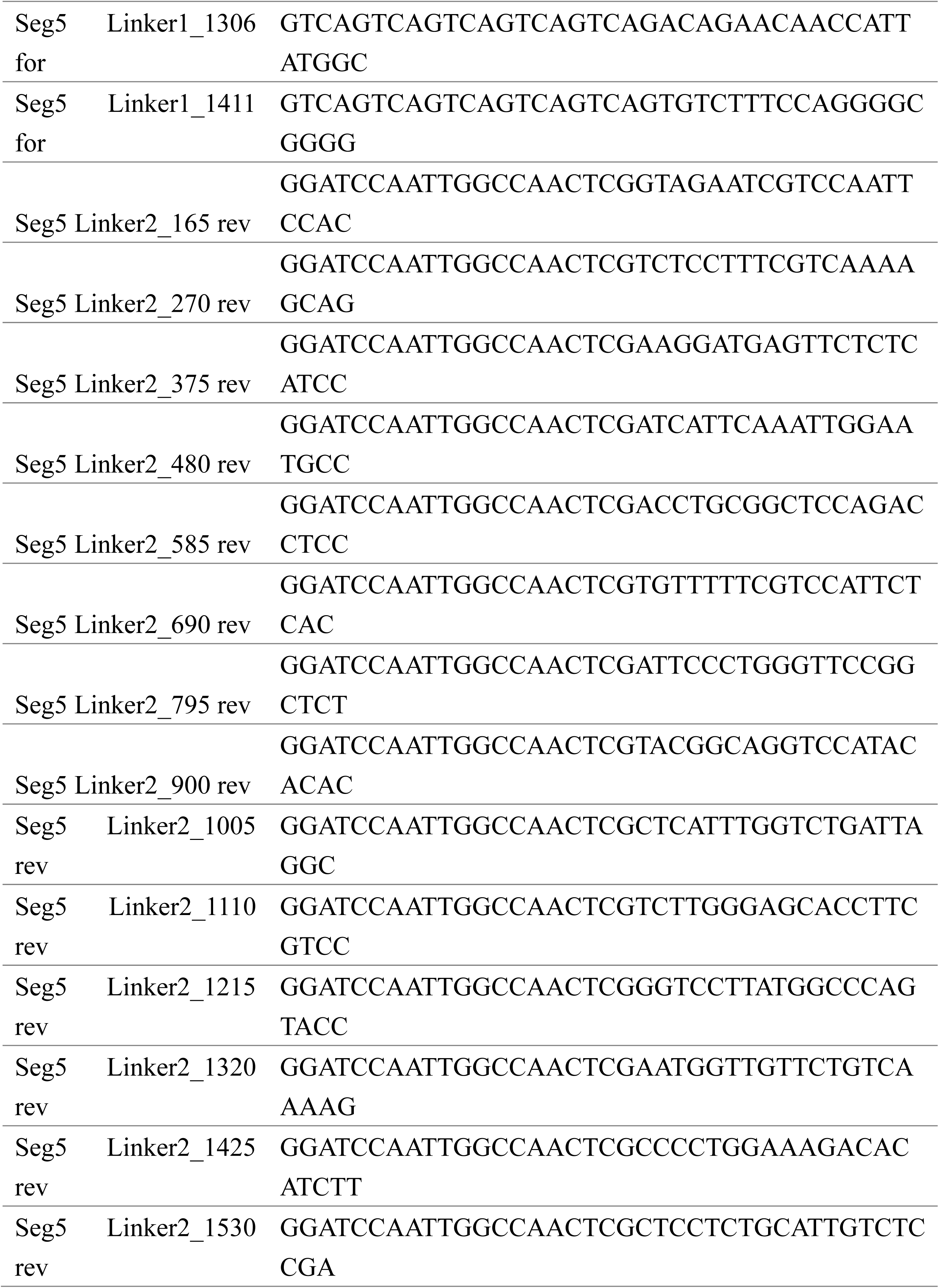
Primers used in this study.

